# Bioinspired living nerve conduit with coupled physiological signals enables functional recovery of peripheral nerve

**DOI:** 10.64898/2026.11.10.765821

**Authors:** Yiqi Cao, Qilong Zhao, Mingxing Peng, Xiufeng Li, Chengbo Liu, Guicai Li, Yumin Yang, Min Wang, Xuemin Du

## Abstract

Long-distance peripheral nerve injury causes severe tissue dysfunction. However, current repair strategies fail to recapitulate the native regenerative microenvironment with coupled physiological signals, hindering functional restoration. Here, we report a bioinspired living neural conduit (LNC) featuring a tri-layered, nerve-like architecture comprising a hydrogel bilayer, nano-grooved ferroelectric polymer stripes, and living Schwann cells, imparting coupled biomechanical, bioelectrical, and biochemical signals. The LNC achieves exceptional adaptability at the injured nerve interface, enabling nerve tissue-matched electrical conductivity, conformal and sutureless integration, and sustained secretion of pro-regenerative exosomes and growth factors (2.2-fold neurotrophic and 1.4-fold angiogenic factor upregulation). In rodent long-gap nerve defect models, the LNC orchestrates intraneural angiogenesis, accelerates axonal outgrowth, and enhances remyelination, achieving unprecedented motor function recovery without neuroma formation or adverse immune responses. By faithfully recapitulating the coupled biophysical and biochemical microenvironment of native nerve repair, the LNC offers a promising strategy towards functional restoration in challenging tissue injuries.

## Introduction

Peripheral nerve injury (PNI) presents a major global healthcare burden, afflicting over 5 million individuals annually due to trauma, disease, and surgical interventions, frequently resulting in long-term motor and sensory deficits^1^. PNI induces a series of pathophysiological events, such as axonal collapse, bioelectrical signal blockades, inner blood vessel disruption, and progressive neuromuscular degeneration, disrupting the microenvironment at the injured nerve and causing great difficulty in the nerve repair, especially for long-distance peripheral nerve defects (> 1 cm in rat)^2^. Currently, autograft remains the clinical gold standard; however, satisfactory functional recovery is achieved in only ∼50% of patients, and this treatment is severely restricted by donor source limitations, structural mismatch between the injured nerve and donor nerve, and surgical complications^3, 4^. As alternatives, artificial nerve grafts (*i.e*., nerve conduits) based on emerging materials have been developed to foster a pro-regenerative microenvironment^5, 6^. Explored nerve conduits usually incorporate anisotropic architectures, electroactive materials, or bioactive cargoes such as exosomes or growth factors, offering a biomechanical, bioelectrical, and/or biochemical active microenvironment to promote cell growth and enhance nerve repair^7–12^. Despite extensive advances, existing nerve conduits still lack the capability of providing synergistic and controllable biomechanical, bioelectrical, and biochemical cues, resulting in unsatisfactory treatment outcomes. In addition, both suturing-induced traumatic neuroma and absence of microvascularization further compromise nerve regeneration and functional recovery^13, 14^. Overcoming these challenges necessitates a paradigm shift toward biomimetic nerve conduits with nerve regeneration-like biomechanical-bioelectrical-biochemical coupling, thus recapitulating natural physiological process including sequential stages of phagocytic debris clearance, vascular endothelial growth factor (VEGF)-mediated in-nerve angiogenesis, guided axonal extension along Bands of Büngner, and ultimately selective synaptogenesis with functional reinnervation of target musculature^15–24^.

Here, we report a bioinspired living nerve conduit (LNC) that couples nerve-like biomechanical, bioelectrical, and biochemical signals to orchestrate immune modulation, intraneural angiogenesis, guided axonal extension, and neuromuscular reconnection, enabling functional reinnervation of target musculature (Figure 1a). Distinct from autografts and conventional nerve conduits, the LNC features a tri-layered, nerve-like architecture comprising (Figure 1a): (i) a natural polymer hydrogel bilayer for conformal and sutureless integration via self-rolling and interfacial adhesion behaviors; (ii) a ferroelectric layer of nano-grooved polydopamine/poly(vinylidene fluoride-*co*-trifluoroethylene) (PDA/P(VDF-TrFE)) stripes for generating biomechanical and bioelectrical cues; and (iii) a living SCs layer for sustained secretion of biochemical cues, including bioactive exosomes, neurotrophic factors, and angiogenic factors. Leveraging the superior functions and coupled physiological signals, the LNC achieves exceptional adaptability at the injured nerve interface, enabling nerve tissue-matched electrical conductivity (7.5 ± 0.9 mS·cm^-1^), robust conformal and sutureless integration (shear strength: ∼80 kPa), and enhanced secretion of pro-regenerative exosomes (3-fold increase; upregulated 42 types of miRNAs related to neural regeneration) and growth factors (2.2- and 1.4-fold enhancements in neurotrophic and angiogenic factors, respectively). In a rodent model of long-distance sciatic nerve amputation, the LNC effectively coordinates early in-nerve angiogenesis, accelerated axonal sprouting, and robust remyelination, enabling unprecedented recovery of in-nerve blood supply and motor function (SFI: -35.7 ± 1.9) without suture-induced trauma, tissue compression, neuroma formation, or detectable, adverse immune responses. By faithfully recapitulating the coupled biophysical and biochemical microenvironment of native nerve tissue regeneration, the LNC offers a powerful translational strategy for repairing long-gap nerve defects, paving a new avenue for regenerative medicine and biohybrid brain-machine interfaces.

**Figure 1.**
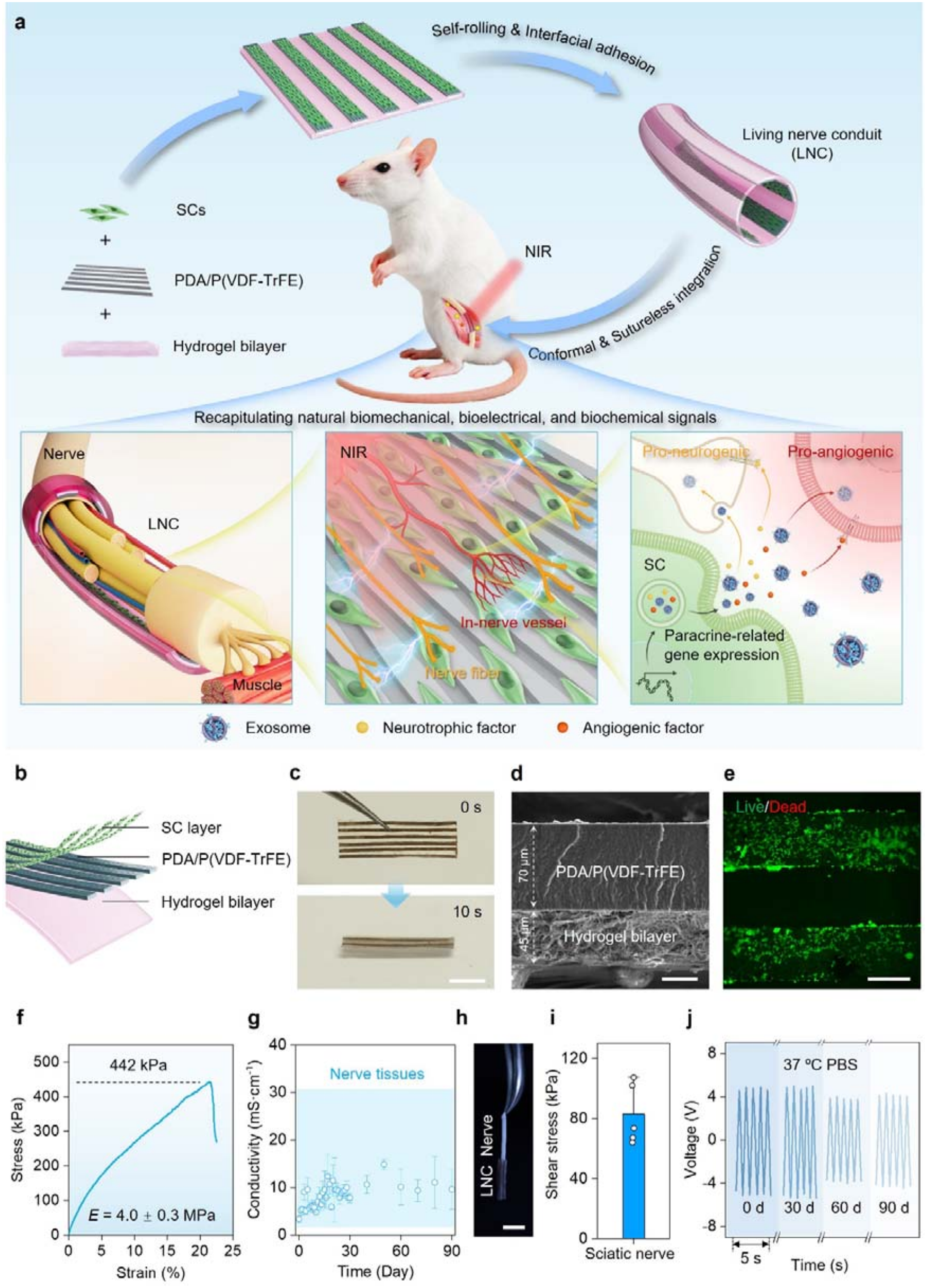
Design and fabrication of the bioinspired LNC. (**a**) Schematic illustration of the LNC, comprising a hydrogel bilayer substrate, nano-grooved PDA/P(VDF-TrFE) ferroelectric polymer stripes, and a living SCs layer. The LNC enables conformal and sutureless integration with amputated sciatic nerves and subsequently promotes revascularization and functional reinnervation, mediated by coupled biomechanical, bioelectrical, and biochemical signals. (**b**) Layout of the tri-layered LNC architecture. Created in BioRender. Siat, I. (2026) https://BioRender.com/xy9i3vz and https://BioRender.com/vo2btzb. (**c**) Time-lapse images illustrating the autonomous self-rolling of the planar LNC into a 3D tubular structure upon immersion in water. Scale bar: 2 mm. (**d**) Cross-sectional scanning electronic image of the LNC, showing a ∼70-μm-thick PDA/P(VDF-TrFE) stripe layer and a ∼45-μm-thick hydrogel bilayer. Scale bar: 30 μm. (**e**) Fluorescence image of LNC, where the incorporated SCs are stained with Calcein AM and Ethd-1 to indicate live and dead cells, respectively. Scale bar: 200 μm. (**f**) Representative strain-stress curve for the LNC under tension. (**g**) Statistics of conductivity of LNC when immersed in 37 °C, phosphate saline buffer (PBS) within 90 days. (**h**, **i**) Interfacial adhesion of LNC onto a segment of *ex vivo* sciatic nerve (**h**), with an average interfacial shear stress of about 80 kPa (**i**). Scale bar: 2 mm. N = 5. (**j**) Open-circuit voltage generated by the LNC at the initial status or immersed in 37 °C, phosphate saline buffer (PBS) within 12 weeks.

## Results

### Design and fabrication of LNC

Recapitulating the complex, multifactorial microenvironment of native neural regeneration remains a fundamental challenge in neural tissue engineering. To address this, we designed a novel LNC featuring a tri-layered, nerve-like architecture that couples biomechanical, bioelectrical, and biochemical signals within a single platform (Figure 1a, b). The outer hydrogel bilayer acts as a biomimetic epineurium that combines nerve tissue-matched mechanics and electrical conductivity with self-rolling and adhesion functions, enabling conformal and sutureless tissue integration to prevent suture-induced trauma. Within this protective shell, an alternating array of nano-grooved ferroelectric polymer stripes and living SCs mimics the anisotropic myelin sheath, synergistically delivering controllable physiological signals. This bioinspired design will endow the LNC with unprecedented structural and functional adaptability to native peripheral nerves^25^.

Following this design strategy, we fabricated the bioinspired LNC via a multistep process (Figure S1, see details in Methods). Briefly, a natural polymer hydrogel bilayer was prepared, consisting of a chitosan hydrogel base and an upper functional layer of Ca^2+^-crosslinked, *N*-hydroxysuccinimide-functionalized alginate (Alg-NHS), and bio-ionic liquid-conjugated polyacrylic acid networks (PAAc-IL). Pre-fabricated nano-grooved PDA/P(VDF-TrFE) ferroelectric polymer stripes were subsequently integrated onto the inner hydrogel surface via micro-mechanical interlocking, followed by seeding with allogenic rat-derived SCs to minimize immunogenicity. As shown in Figure 1c-e, the resulting LNC features a distinct tri-layered architecture: an outer hydrogel bilayer, an inner alternating array of ferroelectric stripes with uniform anisotropic nano-grooves, and a top layer of living SCs exhibiting exceptionally high viability (>98%)^26–28^. Remarkably, the LNC exhibits nerve tissue-level viscoelasticity with a Young’s modulus of 4.0 ± 0.3 MPa, thus minimizing mechanical compression upon implantation (Figure 1f)^29^. Furthermore, the integration of PAAc-IL endows the hydrogel network with nerve tissue-matched ionic conductivity (7.5 ± 0.9 mS·cm^-1^, long-term stability over ∼90 days, Figure 1g, Figure S2), a range optimized to promote electrotaxis while avoiding the cytotoxic or signal-blocking side effects reported at extreme conductivities (> 32 mS·cm^-1^ or < 0.2 mS·cm^-1^)^30–32^. Driven by differential swelling behaviors between the chitosan and Alg-NHS/PAAc-IL layers, the planar LNC autonomously self-rolls into a 3D tubular structure upon hydration, rapidly achieving a curvature of ∼1.0 mm^-1^ within 25 seconds (Figure S3a, b, and Video 1). Such dynamic shape-morphing capability enables conformal, circumferential wrapping around target nerves, thereby resolving the long-standing geometric mismatch associated with autografts (Figure 1c, and Figure S3c)^19, 28^. Concurrently, the LNC establishes robust interfacial bio-adhesion (shear strength: ∼80 kPa) with the neural tissue via synergistic hydrogen bonding and NHS-ester-mediated covalent coupling with tissue surface amines, eliminating suture-induced trauma together with the afterwards neuroma formation (Figure 1h, i, Figure S3c, and Video 2)^33–35^.

Beyond its bioinspired architecture, the LNC is capable of generating nerve-like bioelectrical signals. Upon exposure to NIR irradiation, PDA-mediated photothermal heating triggers the reorientation of P(VDF-TrFE) dipoles within the PDA/P(VDF-TrFE) stripes, inducing reversible polarization changes that establish endogenous-like electric fields (Figure S4a-f)^36, 37^. Specifically, incorporating 1.0 wt% PDA enables the ferroelectric polymer stripes to generate a robust electrical field with a high peak-to-peak open-circuit voltage of ∼ 14 V, which can be facilely tuned by adjusting the PDA concentration or NIR power density (Figure S4c, f). Notably, the NIR-induced electric field exhibits superior long-term stability even under physiological conditions for up to ∼90 days (Figure 1j). Patch-clamp recordings under physiological conditions further confirmed the bioelectric output, which revealed transient, nanoampere-level peak currents dominated by capacitive charging with negligible Faradaic currents, suggesting the safety and efficacy of the ferroelectric layer for cellular electrical stimulation (Figure S4g, h)^37, 38^. Remarkably, unlike conventional electrostimulation approaches that rely on percutaneous wires, bulky energy harvesters, or electrochemical power sources, the LNC wirelessly generates physiological electric fields in a battery-free and non-contact manner, effectively mitigating risks of systemic infection and toxic electrolyte leakage^7, 8, 39–43^. Collectively, these results validate that the bioinspired LNC synergistically integrates nerve-like architecture with precisely tunable biophysical cues, potentially acting as a versatile platform to regulate the behaviors of encapsulated living cells.

### Modulation of SCs within LNC

We next investigate the SC behaviors within the LNC under the modulation by the biomimetic biomechanical and bioelectrical cues (Figure 2a). Under the guidance by the surface anisotropic nano-grooved array, SCs within the LNC exhibit elongated shapes parallel to the oriented nano-grooves, driven by polarized traction stresses along the nano-grooves owing to oriented cytoskeleton fiber (Figure 2b)^44^. The orientation angles of the SCs on such surfaces located within the range of 0-10°, resembling the structure of bands of Büngner for natural nerves (Figure 2c)^45^. The guiding effects between the nano-grooved and the micro-grooved arrays on the SC orientation have been compared, indicating similar cellular elongation, which may upregulate the expression of genes related to myelination (Figure 2b, c)^10^. However, SCs on the control samples with flat surfaces are randomly distributed, which are distinctively different from the natural conditions with anisotropic cellular arrangements (Figure 2b, c)^46^. In addition to the effect of contact guidance for inducing SCs to behave nerve-like oriented arrangements, the biomimetic biomechanical cues can further promote SC migration, exhibiting about 2-fold increase in the migration velocity as compared to the flat controls (Figure 2d, e and Figure S5). Such cellular responses mediated by the biomechanical cues of LNC are relevant to not only the maturation of SCs but also their gene and protein expression, highly compliant with the requirements of neural regeneration^45^.

**Figure 2.**
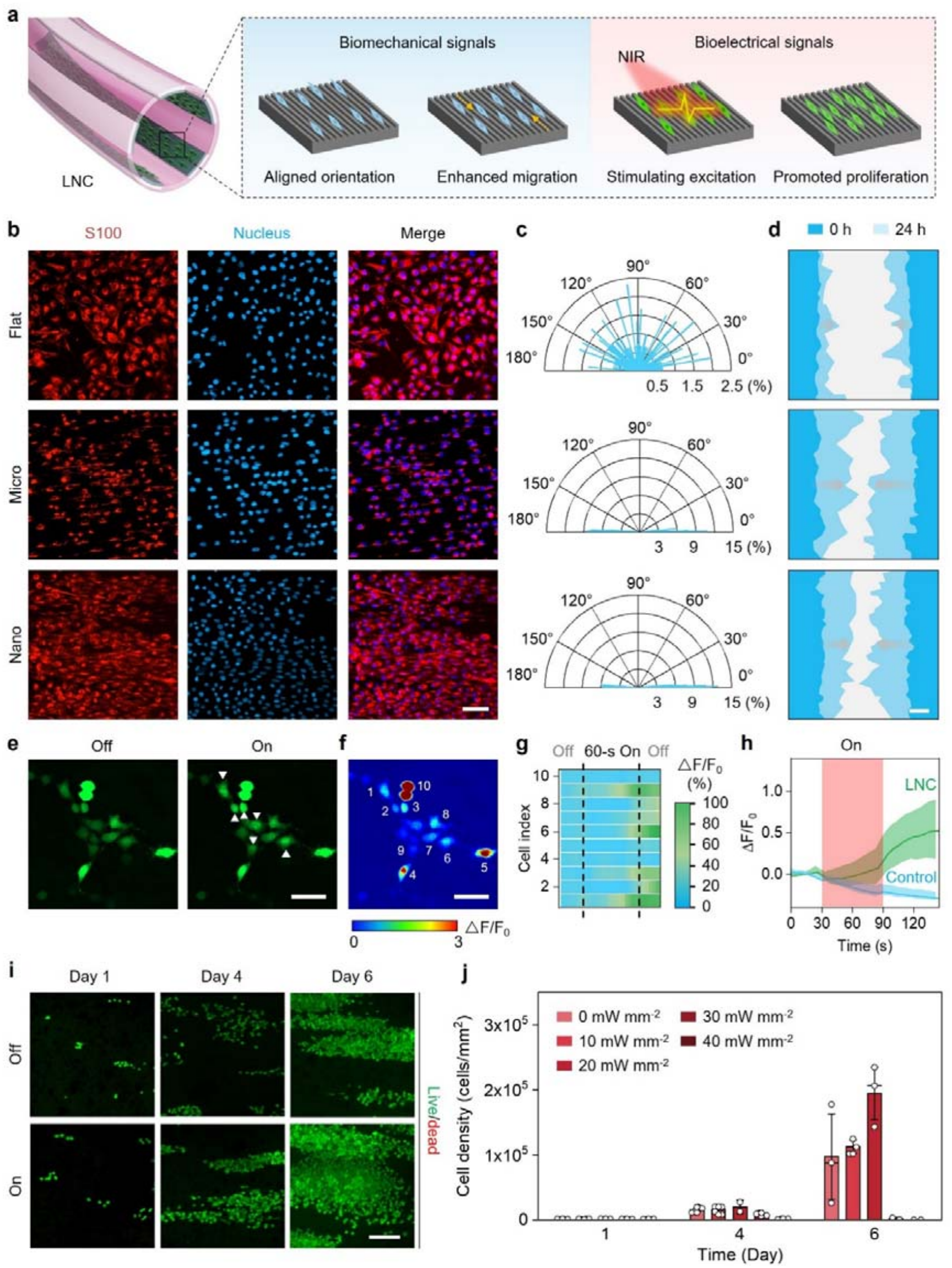
Modulation of SCs within LNC. (**a**) Schematic illustrating the modulation of biomechanical and bioelectrical cues from LNC on the behaviors of the incorporated SCs. (**b**) Representative fluorescence images of SCs with immunochemical staining of S100 and nucleus on different types of samples, including flat PDA/P(VDF-TrFE) film (Flat), PDA/P(VDF-TrFE) films with micro-grooved array on the surfaces (Micro), and PDA/P(VDF-TrFE) films with nano-grooved array on the surfaces (Nano). Scale bar: 50 μm. (**c**) Statistics of cell orientation angles for the SCs. (**d**) Traces of cell migration for the SCs. Scale bar: 200 μm. (**e**) Representative fluorescence images for the SCs stained with fluo4 AM and cultured on the PDA/P(VDF-TrFE) films, before and after NIR irradiation (power density: 20 mW·mm^-2^, 1 Hz frequency, 50% duty factor; recording: 30-s off, 60-s on, and another 30-s off). With white triangles pointing out the cells with obvious fluorescence alternations after NIR irradiation. Scale bar: 50 μm. (**f**) Heat map illustrating the normalized fluorescence changes (ΔF/F_0_) over time for the SCs on the PDA/P(VDF-TrFE) films under NIR irradiation. Scale bar: 50 μm. (**g**) Statistics for the ΔF/F_0_ over time for each SC on the PDA/P(VDF-TrFE) films under NIR irradiation. (**h**) Average ΔF/F_0_ over time for the SCs on the PDA/P(VDF-TrFE) films or the control film (i.e., PDA/poly(vinyl chloride) films with the nano-groove arrays on the surfaces, free of bioelectrical signals under NIR irradiation) and treated with the identical NIR irradiation (power density: 20 mW·mm^-2^, 1 Hz frequency, 50% duty factor; duration: 60 s). N = 10 cells. (**i**) Representative fluorescence images of SCs, stained with Calcein AM and EthD-1, on the PDA/P(VDF-TrFE) films and treated with NIR irradiation at different power densities (0, 10, 20, 30, and 40 mW·mm^-2^) for 6 days. Scale bar: 500 μm. (**j**) Statistics of cell densities for the SCs with or without NIR irradiation. N = 4 separate experiments.

In addition to biomechanical cues, bioelectrical cues are also key signals mediating cellular behaviors during native neural regeneration, where the endogenous electrical fields can switch ion channels and then evoke Ca^2+^ signaling, subsequently regulating a series of SC behaviors related to neural regeneration such as proliferation^47^. To evaluate whether the biomimetic bioelectrical cues of LNC can attain similar effects, intracellular Ca^2+^ imaging for the SCs within LNC is performed. The results reveal that the NIR-mediated bioelectrical cues of LNC can reliably elicit Ca^2+^ influx, as revealed by significantly elevated fluorescence, emulating the physiological Ca^2+^ signaling (Figure 2f and Video 3)^47^. However, control samples absent of bioelectrical cues evoke few SC responses under the identical NIR conditions (Figure 2g, Figure S6a-c, and Video 4). Accordingly, proliferation of SCs within LNC can also be effectively promoted by the NIR-mediated bioelectrical cues, demonstrating a positive correlation to the NIR power densities within a range from 0 to 20 mW·mm^-2^ (Figure 2j). Noted that the density of SCs within LNC under NIR irradiation at the optimized power density of 20 mW·mm^-2^ and after 6 days of culturing is more than doubled as compared to the non-irradiated control, which can accelerate the formation of bands of Büngner and thereby myelination; yet further increase the power density of NIR light leads to apoptotic side effects (Figure 2j and Figure S6d)^47^. Collectively, the biomechanical and bioelectrical cues endowed by LNC enable modulation of SC behaviors including polarization, migration, and proliferation on demand, which not only comply with the requirements of native neural regeneration but also potentially modulate the paracrine activity of SCs via activation of specific signaling pathways^45, 48^.

### Secretion of exosomes and growth factors

Exosomes and growth factors, typically nerve growth factor and vascular endothelial growth factor secreted by endogenous SCs, are essential mediators for biochemical signaling in peripheral nerve regeneration^48–50^. To verify the pro-regenerative potential, we evaluate the secretion of exosome and growth factors by LNC with incorporated SCs (Figure 3a). Transmission electron imaging and western blotting analyses into the characteristic membrane proteins both indicate that LNC is capable of secreting massive exosomes with typical cup-shaped nanostructures (Figure S7a, b). Nanoparticle tracking analysis (NTA) further quantifies the contents of exosomes secreted by LNC with the nano-grooved array surfaces, indicating ∼ 1×10^9^ particles·mL^-1^ in total and corresponding to a per cell level of 551 ± 53 particles, which is about 26% upregulation as compared to the per cell value for control samples with flat surfaces and hence lack of biomimetic biomechanical cues (Figure 3b and Figure S7c-h). Interestingly, despite similar modulating effects on SC behaviors as indicated above, the samples with micro-grooved array surfaces cannot enhance exosome secretion for the incorporated SCs like LNC (Figure 3b). On the contrary, the exosome secretion is inhibited, implying distinct effects between these similar biomechanical cues. In addition to the biomimetic biomechanical cues, the regulation effects of the biomimetic bioelectrical cues on exosome secretion are also assessed via NTA quantification. The results demonstrate that the biomimetic bioelectrical cues of LNC endowed by the NIR-induced electrical fields can further upregulate exosome section, realizing almost 3-folds increases in secreting contents per cell as compared to the non-irradiated conditions (Figure 3c and Figure S7). However, non-ferroelectric samples involving only photothermal effects under the identical NIR irradiation conditions exhibit a comparable exosome-secreting level to the non-irradiated conditions, suggesting essential role of the bioelectrical signals in enhancing exosome secretion (Figure 3c and Figure S7). LNC with such enhanced exosome secreting capabilities will attain superior biochemical signaling over conventional SC transplantation strategies^51^.

**Figure 3.**
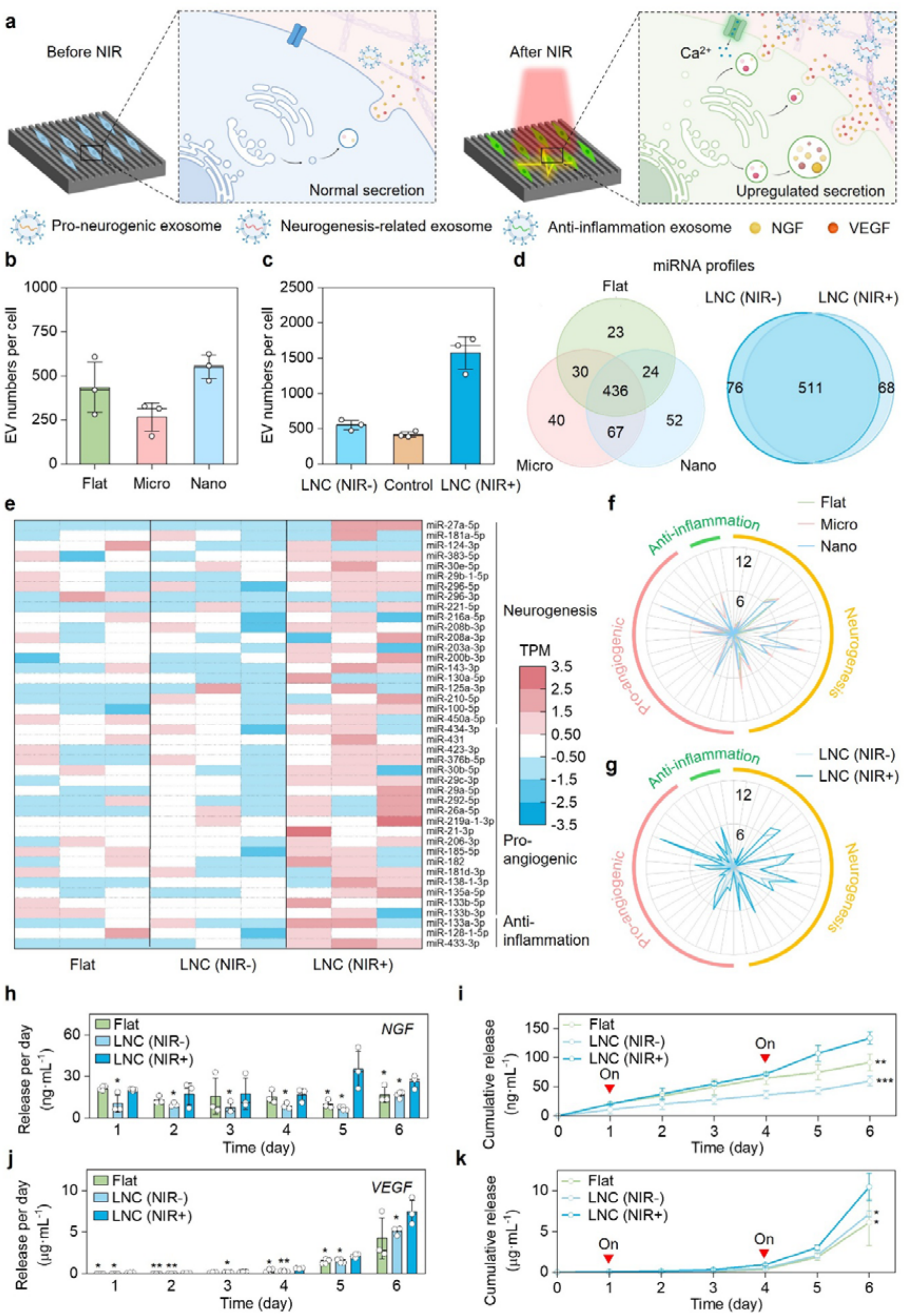
Secretion of exosomes and growth factors. (**a**) Schematic illustrating the modulation of biomechanical and bioelectrical cues from LNC on their exosome and growth factor secretion. Created in BioRender. Siat, I. (2026) https://BioRender.com/icrl8ko. (**b**) Statistics of extracellular vehicle (EV) number secreted per cell for the SCs cultured on different types of samples, including flat PDA/P(VDF-TrFE) film (Flat), PDA/P(VDF-TrFE) films with micro-groove array microstructures on the surfaces (Micro), and PDA/P(VDF-TrFE) films with nano-groove array nanostructures on the surfaces (Nano, i.e., the ferroelectric layer of the LNC). N = 3 separate experiments. (**c**) Statistics of EV number secreted per cell for the SCs on the control film (i.e., PDA/poly(vinyl chloride) films with the nano-groove arrays on the surfaces, free of bioelectrical signals under NIR irradiation) under NIR irradiation (power density: 20 mW·mm^-2^, 1 Hz frequency, 50% duty factor) or within the LNC with and without NIR irradiation (LNC (NIR+) and LNC (NIR-)). N = 3 separate experiments. (**d**) Venn diagrams for the variances in the miRNA profiles among the groups with different types of biomechanical or bioelectrical cues. (**e**) Expression of miRNAs related to neurogenesis, pro-angiogenic, and anti-inflammation functionalities in the exosomes secreted by the SCs among different groups. N = 3 separate experiments. (**f**, **g**) Comparison into the average expression of miRNAs related to neurogenesis, pro-angiogenic, and anti-inflammation functionalities among the groups with different types of biomechanical (**f**) or bioelectrical cues (**g**). (**h-k**) Release profiles of NGF (**h**, **i**) and VEGF (**j**, **k**) by the SCs within different groups with different types of biomechanical or bioelectrical cues. Release per day for NGF: Day 1: LNC (NIR+) vs Flat: P = 0.216, LNC (NIR+) vs LNC (NIR-): P = 0. 039, Day 2: LNC (NIR+) vs Flat: P = 0.230, LNC (NIR+) vs LNC (NIR-): P = 0.011, Day 3: LNC (NIR+) vs Flat: P = 0.437, LNC (NIR+) vs LNC (NIR-): P = 0.013, Day 4: LNC (NIR+) vs Flat: P = 0.363, LNC (NIR+) vs LNC (NIR-): P = 0.029, Day 5: LNC (NIR+) vs Flat: P = 0.022, LNC (NIR+) vs LNC (NIR-): P = 0.015, Day 6: LNC (NIR+) vs Flat: P = 0.045, LNC (NIR+) vs LNC (NIR-): P = 0.017, N = 3 separate experiments; cumulative release for NGF: LNC (NIR+) vs Flat: P = 0.008, LNC (NIR+) vs LNC (NIR-): P < 0.001, N = 3 separate experiments; Release per day for VEGF: Day 1: LNC (NIR+) vs Flat: P = 0.017, LNC (NIR+) vs LNC (NIR-): P = 0. 017, Day 2: LNC (NIR+) vs Flat: P = 0.002, LNC (NIR+) vs LNC (NIR-): P = 0.004, Day 3: LNC (NIR+) vs Flat: P = 0.120, LNC (NIR+) vs LNC (NIR-): P = 0.030, Day 4: LNC (NIR+) vs Flat: P = 0.036, LNC (NIR+) vs LNC (NIR-): P = 0.002, Day 5: LNC (NIR+) vs Flat: P = 0.027, LNC (NIR+) vs LNC (NIR-): P = 0.022, Day 6: LNC (NIR+) vs Flat: P = 0.082, LNC (NIR+) vs LNC (NIR-): P = 0.036, N = 3 separate experiments; cumulative release for VEGF: LNC (NIR+) vs Flat: P = 0.008, LNC (NIR+) vs LNC (NIR-): P < 0.001, N = 3 separate experiments. Statistical analyses into the data shown in **h**, **i**, **j**, and **k** were performed using one-way ANOVA with two-tailed Student’s *t*-test.

Considering profiles of microRNA (miRNA) cargoes within the exosomes determine their specific bio-functionalities, we then perform bulk miRNA sequencing for the exosomes among samples with different biomechanical and bioelectrical microenvironments. The results indicate that the variations in biomechanical and bioelectrical cues result in obvious differences in miRNA expression (Figure 3d). Notably, a variety of miRNAs related to neural regeneration, including 20 neurogenesis-related genes, 19 pro-angiogenic genes, and 3 anti-inflammation genes, are upregulated expressed in the exosomes secreted by LNC with the biomimetic biomechanical and bioelectrical cues, which will be advantageous in promoting neural regeneration over the control samples without NIR-induced electrical fields or lacking the nano-grooved array surfaces (Figure 3e-g). However, the control samples with the micro-grooved array surfaces cannot attain significant upregulation in the expressions of these neural regeneration-related miRNAs under the identical NIR conditions (Figure S8).

Next, we quantify the secretion of NGF and VEGF from LNC and compare it with the conditions without the biomechanical or bioelectrical cues. Enzyme-linked immunosorbent assay reveals that the samples with biointegrated design can achieve sustained secretion of NGF and VEGF, eliminating concerns related to insufficient payloads and progressive degradation for conventional delivery strategies (Figure 3h-k)^19^. Moreover, LNC exhibits stronger secreting capabilities for both NGF and VEGF than those of flat controls; and strikingly, the bioelectrical modulation via NIR irradiation can further enhance the secretion of the neurotrophic and angiogenic factors by 2.2 and 1.4 folds, respectively (Figure 3i, k). These up-regulated quantities will meet the physiological requirements in the natural peripheral nerve regeneration^52, 53^. Harnessing the concurrent upregulation of exosomes and growth factors with diverse bio-functionalities, the LNC offers physiologically faithful biochemical cues to orchestrate key peripheral nerve repair events including angiogenesis, axonal sprouting, remyelination, and target reinnervation.

### Mechanistic study

We subsequently investigate how the biomimetic biomechanical and biochemical cues of LNC modulate the exosome and growth factor secretion for SCs, which remain unknown according to existing literature. To elucidate the underlying mechanism, bulk RNA sequencing is performed for the SCs within LNC, with or without the NIR-induced bioelectrical modulation. When comparing LNC groups with the control groups where SCs are grown on the samples with flat surfaces, both absences of NIR irradiation, we identify 88 types of genes with significant differences, of which 55 types are upregulated (Figure 4a and Figure S9a). Among these upregulated genes, there are 7 types pertinent to cytoskeleton organization (e.g., *Ccl27*), exocytosis (e.g., *Mip*, *Osm*), angiogenesis (e.g., *Fut1*), and axon extension (e.g., *L1cam*, *Pard6a*, *Cntf*), according to GO enrichment (Figure 4b). Pathway enrichment analysis further points enrichments in processes related to exosome and growth factor secretion (Figure 4c). However, both gene expression and gene ontology analysis indicate that though numerous genes exhibit significant alterations when comparing the groups between the micro-groove array surfaces and the flat controls, among which few upregulated genes are associated with biosynthesis or exocytosis (Figure S10). In contrast, the biomimetic biomechanical cues derived by the nano-groove array surfaces exhibit pronounced effects on biosynthesis or exocytosis via the activation of biomechanic-related signaling pathways, implying possible mechanisms for the upregulation of exosome and growth factor section for LNC^54^.

**Figure 4.**
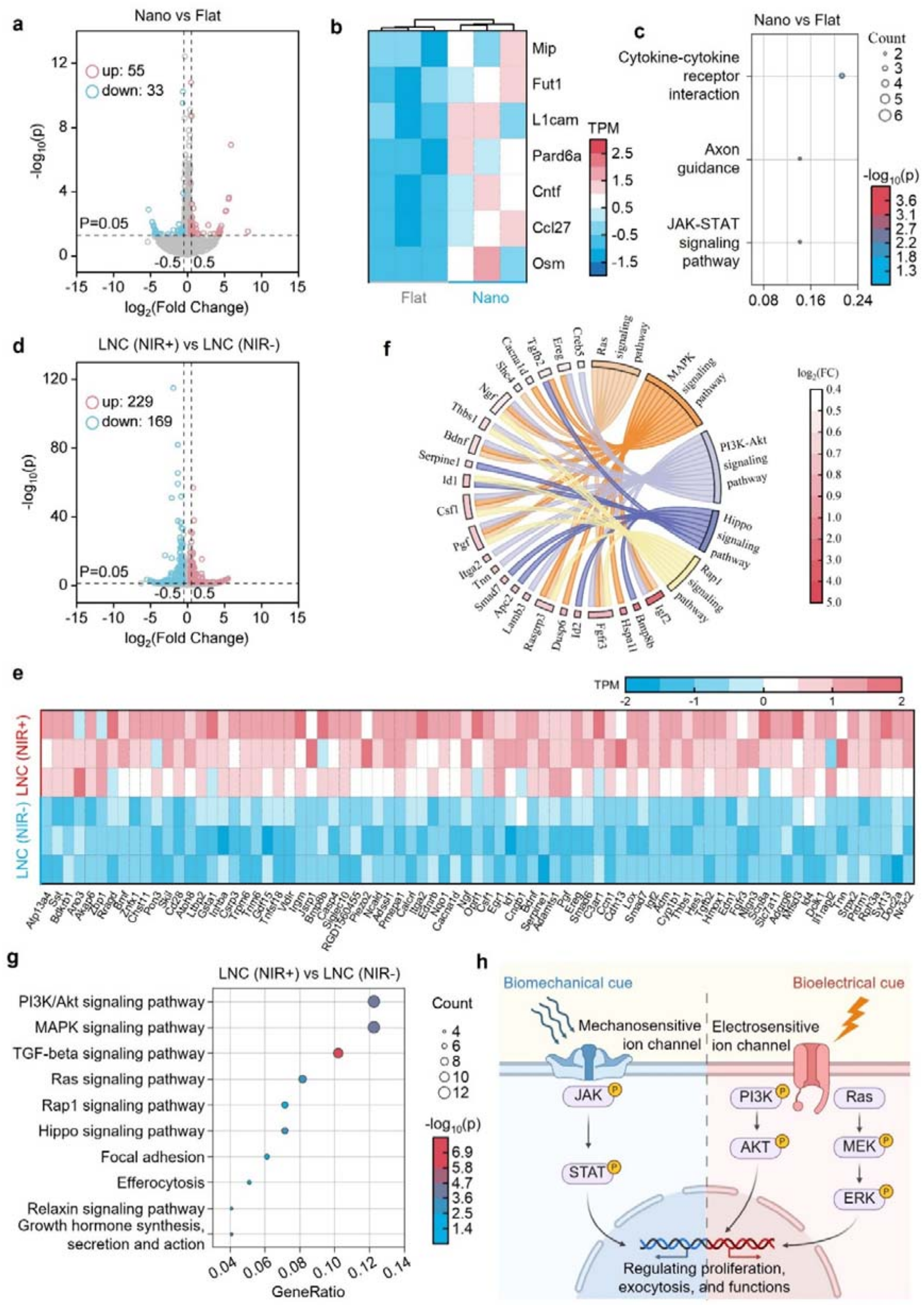
Transcriptomic profiles of SCs within LNC. (**a**) Variances in the RNA expression between the Nano and Flat groups. (**b**) Upregulated expression of genes related to cytoskeleton organization, exocytosis, angiogenesis, and axon extension between the Nano and Flat groups. (**c**) KEGG enrichment for the signaling pathways activated by biomechanical cues. (**d**) Variances in the RNA expression between the LNC (NIR-) and LNC (NIR+) groups. (**e**, **f**) Upregulated expression of genes related to Ca^2+^ signaling, biosynthesis, exocytosis, angiogenesis, axon extension, and neuroinflammation between the LNC (NIR-) and LNC (NIR+) groups. (**g**) KEGG enrichment for the signaling pathways activated by bioelectrical cues. (**h**) Summary of molecular mechanisms for the modulation of SCs by the biomechanical and bioelectrical signals. Created in BioRender. Siat, I. (2026) https://BioRender.com/i0czkq0.

When comparing the conditions with and without NIR irradiation, we observe a massive number of genes exhibit significantly different expression, among which the amounts of upregulated genes reaching 229, inferring more significant effects on SC regulation for the biomimetic bioelectrical cues than those of biomechanical cues (Figure 4d and Figure S9b). Importantly, there are 79 types of upregulated genes engaged in Ca^2+^ signaling (e.g., *Atp13a4*, *Tgfb2*, *Bdkrb1*), biosynthesis (e.g., *Egr1*, *Igf2*, *C3ar1*), exocytosis (e.g., *Rph3a*, *Syt13*, *Doc2a*), angiogenesis (e.g., *Tnn*, *Srpx2*, *Pgf*), axon extension (e.g., *Nlgn3*, *Edn1*, *Ngf*), and neuroinflammation (e.g., *Hes1*, *Prl7a3*, *Cyp1b1*), which will determine exosome and growth factor secretion, as well as miRNA expression (Figure 4e)^55^. Gene ontology analysis of all significantly differentially expressed genes reveals that various biophysics-related signaling pathways, such as phosphatidylinositol 3-kinase (PI3K)/Akt, mitogen-activated protein kinase (MAPK), Ras signaling pathways, etc., are enriched and the most counted (Figure 4f, g and Figure S11). Together, the biomimetic biomechanical and bioelectrical cues enabled by LNC can activate specific signaling pathways for the incorporated SCs, therefore modulating their behaviors and upregulating the secretion of exosomes and growth factors with anti-inflammation, pro-angiogenic, and neurogenesis-related functions, which will potentially promote vascularization-coupled neural regeneration.

### Synergistic angiogenesis and neural regeneration

To assess pro-regenerative capabilities of LNC, we first perform transwell experiments through coculturing LNC respectively with rat-derived macrophages, endothelial cells (ECs), and model neuronal cells (rat phaeochromocytoma (PC12) cells) for evaluating its effects on macrophage polarization, vascular tubule formation and neurite outgrowth, which are hallmarks for immune regulation, vascularization, and neural regeneration (Figure S12a)^23^. Fluorescence microscopies verify that the SCs within LNC can realize intercellular communications with the co-cultured cells, as evidenced by the uptake of PKH26 labelled exosomes secreted by LNC, while fluorescence signals are absent in the SC-free negative controls (Figure S12b). Based on such biochemical signals, LNC demonstrates robust immune-regulating effects, as evidenced by immunofluorescence of iNOS and CD206 for identifying macrophages with M1 and M2 phenotypes, respectively (Figure S12c). Aligned with the upregulated secretion of anti-inflammatory miRNAs under the NIR-irradiated conditions, the group treated with the LNC and NIR irradiation exhibit the highest ratio in the pro-regenerative M2 phenotypes, significantly higher than those in groups with only photothermal or biomechanical signals (Figure S12d-f). Moreover, owing to the upregulated angiogenic miRNAs and factors under NIR irradiation that regulate post-transcriptional processes and directly modulate target cell behaviors, LNC obviously accelerate the organizations of ECs into vascular tubules with well-defined capillary-like branched networks (Figure S12g). The numbers of circles and nodes in the LNC groups, both key indicators of angiogenesis, are about 50% and 48% higher than those in the groups without NIR irradiation, respectively, validating remarkable pro-angiogenic capabilities of LNC with hyper-physiological signals (Figure S12h, i). Moreover, LNC can further enhance PC12 cell elongation, as revealed by obvious increases in their length-to-width ratios as compared to the negative control groups or the NIR-free groups, implying its superior potential in promoting neurite outgrowth and axon extension (Figure S12j). Immunofluorescence and reverse transcription-quantitative polymerase chain reaction (RT-qPCR) of βIII-tubulin, a typical neuronal marker, both reveal significantly upregulated expressions in the LNC group compared with the SC-free and NIR-free conditions, suggesting pronounced effects of LNC on axonal development and neuronal maturation (Figure S12k, l). These *in vitro* results demonstrate remarkable immune-regulating and pro-regenerative potentials of LNC for anti-inflammation, vascularization, and neural regeneration.

We next implant LNC onto amputated sciatic nerves of rats, with a gap of 1 cm, and then evaluate *in vivo* tissue regeneration processes (Figure 5a). As shown in Figure 5b, LNC can be facilely implanted onto amputated sciatic nerves through water spraying, subsequently realizing conformal contact and sutureless fixation owing to its self-rolling and bio-adhesive characteristics, free of secondary damage to residual nerve tissues caused by sutures commonly required by conventional autografts or neural conduits^35^. After implantation, the LNC is irradiated with 808-nm NIR light every day in a remote-controlled manner, with the penetration depth (10 mm) and temperature rise (∼0.8 K within the entire irradiation process) respectively meeting the transdermal activation and biosafety criteria according to the standards (IEC 60825-1:2014 and ISO 14117:2012, Fig. S13)^56^. Within the critical time window for peripheral nerve repair^2^, the LNC demonstrates durable stability in structure, mechanics, and the capability in generating electrical fields under NIR irradiation, which can eliminate compression to nerve tissues and foreign body responses (with no statistically significant differences in nerve fiber diameters and inflammatory area thickness compared with the contralateral non-operated sites) benefited from its conformal contact and sutureless fixation (Figure 5b and Figure S14). Moreover, the viability of the incorporated SCs can also be well preserved within the whole implantation period (Fig. S15). Collectively, the LNC can therefore provide durable biomechanical, bioelectrical, and biochemical cues for promoting peripheral nerve tissue regeneration.

**Figure 5.**
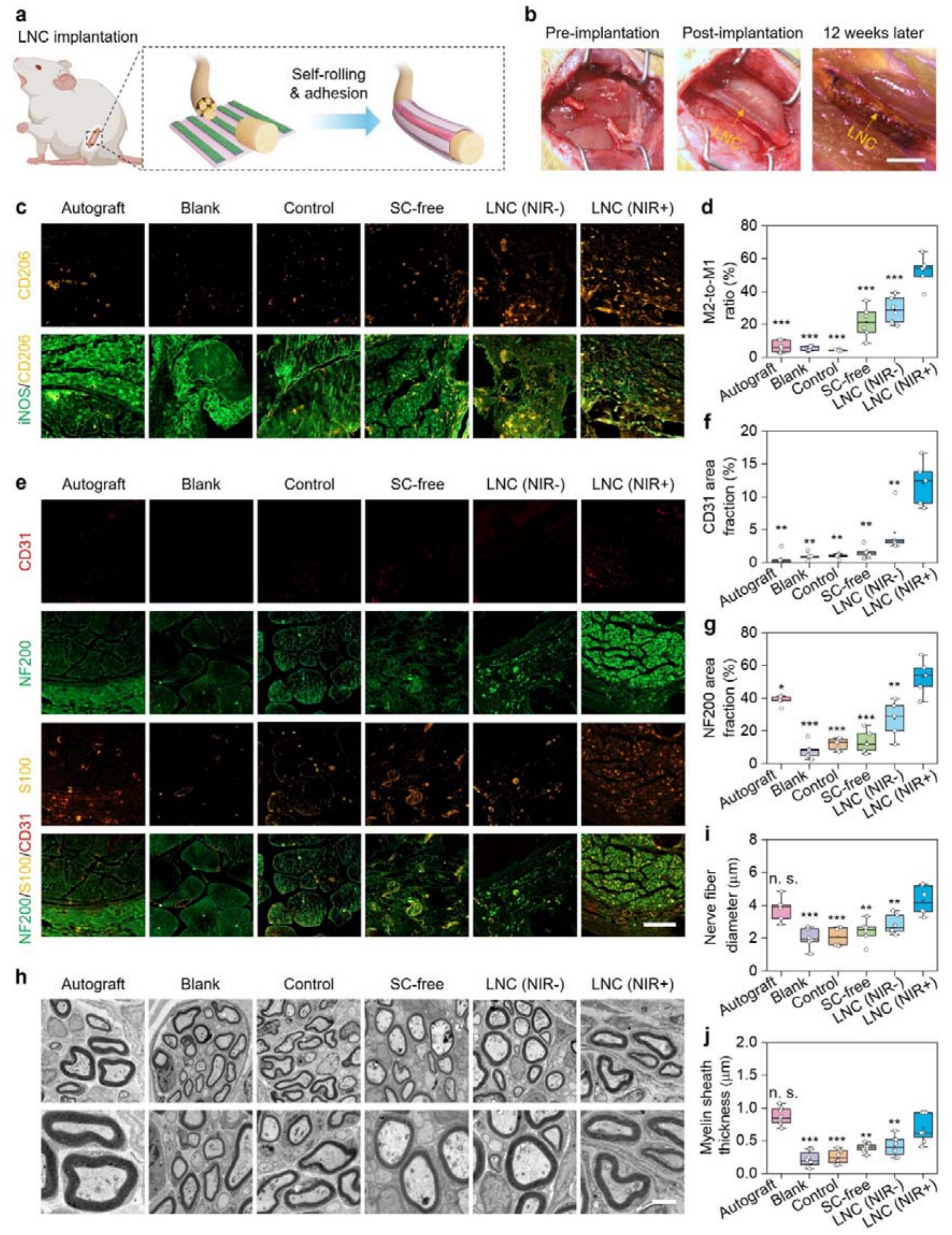
Synergistic angiogenesis and neural regeneration. (**a**) Schematic illustrating the *in vivo* implantation of LNC onto the amputated sciatic nerves of rats. (**b**) Representative pictures for the amputated sciatic nerves of rats before, immediately after LNC implantation, 12 weeks later. Scale bar: 5 mm. (**c**) Representative fluorescence images of regenerated nerve tissues with immunochemical staining of iNOS and CD68, after 12-week implantation of different types of samples: Autograft: a segment of autogenous nerve via suturing fixation. Blank: no additional treatments. Control: SC-free, non-ferroelectric PVC-based samples, followed by post-operative NIR irradiation (power density: 20 mW·mm^-2^, 1 Hz frequency, 50% duty factor, duration: 10 min, performed every 2 days post-surgery). SC-free: SC-free, ferroelectric samples, treated with the NIR irradiation after surgery. LNC (NIR-): LNC without NIR irradiation. LNC (NIR+): LNC with the NIR irradiation after surgery. Scale bar: 100 μm. (**d**) Statistics of M2-to-M1 ratio for the regenerated nerve tissues after 12-week implantation. LNC (NIR+) vs Autograft: P < 0.001, LNC (NIR+) vs Blank: P < 0.001, LNC (NIR+) vs Control: P < 0.001, LNC (NIR+) vs SC-free: P < 0.001, LNC (NIR+) vs LNC (NIR-): P < 0.001. N = 7 separate experiments. (**e**) Representative fluorescence images of regenerated nerve tissues, with immunochemical staining of CD31, NF200, and S100, after 12-week implantation. Scale bar: 100 μm. (**f**, **g**) Statistics of area fractions for CD31 (**f**) and NF200 (**g**) for the regenerated nerve tissues after 12-week implantation. CD31: LNC (NIR+) vs Autograft: P = 0.001, LNC (NIR+) vs Blank: P = 0.002, LNC (NIR+) vs Control: P = 0.002, LNC (NIR+) vs SC-free: P = 0.002, LNC (NIR+) vs LNC (NIR-): P = 0.009. N = 5 separate experiments. NF200: LNC (NIR+) vs Autograft: P = 0.041, LNC (NIR+) vs Blank: P < 0.001, LNC (NIR+) vs Control: P < 0.001, LNC (NIR+) vs SC-free: P < 0.001, LNC (NIR+) vs LNC (NIR-): P = 0.007. N = 5 separate experiments. (**h**) Representative transmission electronic images of regenerated nerve tissues after 12-week implantation. Scale bar: 20 μm. (**i**, **j**) Statistics of nerve fiber diameter (**i**) and myelin sheath thickness (**j**) for the regenerated nerve tissues after 12-week implantation. Nerve fiber diameter: LNC (NIR+) vs Autograft: P = 0.316, LNC (NIR+) vs Blank: P < 0.001, LNC (NIR+) vs Control: P < 0.001, LNC (NIR+) vs SC-free: P = 0.002, LNC (NIR+) vs LNC (NIR-): P = 0.009. N = 8. Myelin sheath thickness: LNC (NIR+) vs Autograft: P = 0.066, LNC (NIR+) vs Blank: P < 0.001, LNC (NIR+) vs Control: P < 0.001, LNC (NIR+) vs SC-free: P = 0.004, LNC (NIR+) vs LNC (NIR-): P = 0.009. N = 8. Statistical analyses into the data shown in **d**, **f**, **g**, **i**, and **j** were performed using one-way ANOVA with two-tailed Student’s *t*-test.

Under the guidance by these LNC-derived signals, injured sciatic nerve tissues exhibit significant regeneration after 12 weeks, as revealed in hematoxylin and eosin (H&E) staining, with the regenerated tissues reconnecting the two ends of the stumps and no signs of neuroma, eliminating voids and the risks of neuroma-induced declines in functional recovery for the “gold-standard” autografts that require suturing fixation (Figure S16)^57^. However, other experimental groups including sham-operated blank controls, inclusion of photothermal effects alone, absence of biochemical cues, and absence of bioelectrical cues demonstrate obviously delayed in situ regeneration (Figure S16). The immunofluorescence of iNOS and CD206 indicates that LNC exhibits minimal macrophage infiltration, meanwhile inducing a higher localization of pro-regenerative M2 phenotypes at the middle of regenerative bridges than that of pro-inflammatory M1 macrophages (Figure 5c). In contrast, macrophages in other experimental groups including the autografting group exhibit increased ratios of M1 to M2 population, which are consistent with the bulk miRNA sequencing that lack the anti-inflammation-related expression (Figure 5d and Figure 3e). To compare the regeneration of nerve tissues including blood vessels, axons, and myelin sheaths among different groups, we then perform immunochemical assessments for the dissected tissues at the middle of regenerative bridges after 12 weeks post-surgeries through staining the EC marker CD31, the mature neuron marker neurofilament-200 (NF200), the SC marker S100 (Figure 5e). The results demonstrate potent efficacy of LNC in promoting neo-nerve tissue formation, with even slightly higher densities of blood vessels, axons, and myelin sheaths than those in the autograft-treated groups, and significantly advantageous over the control groups with only photothermal effects or absence of biomimetic bioelectrical or biochemical cues (Figure 5f, g and Figure S17). It can further be validated by transmission electron microscopy, pointing that the neo-nerve tissues achieve large axonal diameter (4.28 ± 0.96 μm) and thick myelin sheath (0.69 ± 0.18 μm) in the LNC group, comparable to those in the autograft group and superior over other groups (Figure 5h-j). Compared with nerve conduits possessing only photothermal effects, bioelectrical or biochemical inducements, LNC with coupled physiological signals can better facilitate synergistic regeneration of nerve tissues including axon, myelin sheath, and in-nerve vessels, which will assist the reconstruction of neuro-vascular cross-talks and thereby promote functional recovery for injured nerves^23^.

### Recovery of blood supply and motor functions

Functional recovery, especially restoration of blood supply and reinnervation, remains grand challenges in peripheral nerve regeneration for existing strategies^19, 21, 22^. In the rodent models with long-distance sciatic nerve defects, we further evaluate the functional recovery assisted by LNC, which has demonstrated superior capabilities in facilitating vascularization and neural regeneration (Figure 6a). Microcirculation within nerve tissues plays a pivotal role in providing nutrients and maintaining metabolisms for the survival of cells, as well as regulating the homeostasis of other nerve tissue functions^58^. However, to our best knowledge, none of existing studies provide direct evidence to the restoration of blood supply within neo-formed nerve tissues. Considering that LNC can effectively promote the regeneration of vessels, we then examine whether the regenerated vessels can restore functions of blood supply through photoacoustic imaging (Figure 6b). After 12 weeks of implantation, LNC-treated regenerated nerves exhibit apparent signs of blood supply within the tissues, with a blood flowing rate of 0.51 ± 0.13 mm·s^-1^ as measured by the line scanning of the vessels denoted with arrows (Figure 6c, d and Figure S15). Remarkably, the microcirculation function has been recovered to the equivalent level of normal conditions for a rat without peripheral nerve injury, demonstrating a blood flowing rate of 0.66 ± 0.12 mm·s^-1^ for the natural vessels within the intact sciatic nerves (Figure 6c, d and Figure S15). However, even with existences of vascular structure, the vessels within the neo-formed nerve tissues in the control group are occlusive properly hindered by nerve scar, as evidenced by no signs of blood flows (Figure 6c, d and Figure S18). These results provide direct evidence that blood supply within the regenerated nerves can be restored, while absence of blood supply leads to severe problems of central necrosis and finally dysfunction frequently observed in both autografts and artificial nerve grafts^1^, implying potent capabilities of LNC on the functional recovery of injured nerves.

**Figure 6.**
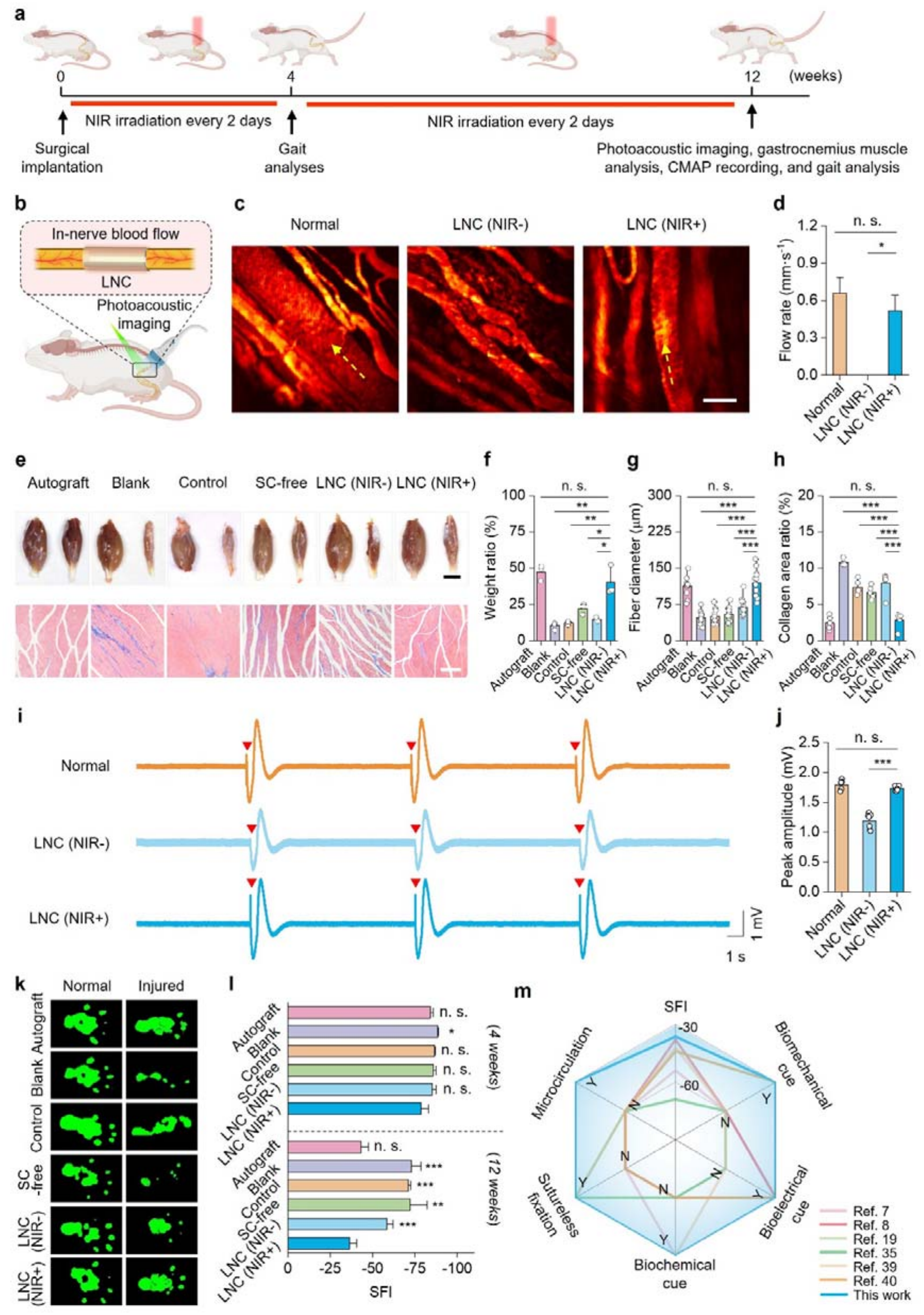
Recovery of blood supply and motor functions. (**a**) Experimental scheme of evaluation into the functional recovery for the rats with amputation of sciatic nerves. (**b**) Schematic illustrating the photoacoustic imaging for assessing the blood flow for the in-nerve vessels. (**c**) Representative photoacoustic images of normal sciatic nerve (Normal) or LNC-implanted (with or without NIR irradiation) regenerated nerves. Dashed arrows illustrate the direction of blood flow. (**d**) Statistics of blood flow rate for the in-nerve vessels. LNC (NIR+) vs Normal: P = 0.504, LNC (NIR+) vs LNC(NIR-): P = 0.047. N = 3. (**e**) Pictures of gastrocnemius muscles collected from both hind limbs (Upper) and microscopic images of gastrocnemius muscles collected from the injured sides with Masson’s staining (Bottom) after 12-week treatments. Scale bar: 100 μm. (**f**-**h**) Statistics of weight ratio for the gastrocnemius muscles between the injured and non-treated sides (**f**), as well as muscle fiber diameter (**g**) and collagen area ratio (**h**) for the gastrocnemius muscles at the injured sides after 12-week treatments. Weight ratio: LNC (NIR+) vs Autograft: P = 0.367, LNC (NIR+) vs Blank: P = 0.008, LNC (NIR+) vs Control: P = 0.010, LNC (NIR+) vs SC-free: P = 0.046, LNC (NIR+) vs LNC (NIR-): P = 0.013. N = 3 separate experiments. Fiber diameter: LNC (NIR+) vs Autograft: P = 0.493, LNC(NIR+) vs Blank: P < 0.001, LNC(NIR+) vs Control: P < 0.001, LNC (NIR+) vs SC-free: P < 0.001, LNC (NIR+) vs LNC (NIR-): P < 0.001. N = 12. Collagen area ratio: LNC (NIR+) vs Autograft: P = 0.603, LNC (NIR+) vs Blank: P < 0.001, LNC(NIR+) vs Control: P < 0.001, LNC (NIR+) vs SC-free: P < 0.001, LNC (NIR+) vs LNC (NIR-): P < 0.001. N = 5 separate experiments. (**i**) CMAP traces of normal sciatic nerves and regenerated nerves treated by LNC with or without NIR irradiation. (**j**) Statistics of CMAP peak amplitude. LNC (NIR+) vs Normal: P = 0.137, LNC (NIR+) vs LNC (NIR-): P < 0.001. N = 3 separate experiments. (**k**) Representative foot traces for the rats after 12-week treatments. (**l**) Statistics of SFI for the rats after 4- and 12-week treatments. 4 weeks: LNC (NIR+) vs Autograft: P = 0.163, LNC (NIR+) vs Blank: P = 0.040, LNC(NIR+) vs Control: P = 0.069, LNC (NIR+) vs SC-free: P = 0.084, LNC (NIR+) vs LNC (NIR-): P = 0.102. N = 4 separate experiments. 12 weeks; LNC (NIR+) vs Autograft: P = 0.059, LNC (NIR+) vs Blank: P < 0.001, LNC(NIR+) vs Control: P < 0.001, LNC (NIR+) vs SC-free: P = 0.007, LNC (NIR+) vs LNC (NIR-): P < 0.001. N = 4 separate experiments. (**m**) Comparation LNC with existing strategies for PNI^7, 8, 19, 35, 39, 40^, in terms of biomimetic characteristics, biosafety, and functional recovery. Y: Yes, N: No. Statistical analyses into the data shown in **d**, **f**, **g**, **h**, **j**, and **l** were performed using one-way ANOVA with two-tailed Student’s *t*-test.

Finally, the recovery of motor functions for hind limbs is evaluated. Motor dysfunction ascribed to innerved muscle atrophy is one of the most predominant causes of peripheral nerve injuries to disability^23^. Accordingly, we analyze the atrophy of gastrocnemius muscles, which are innerved by sciatic nerves and can represent degrees of functional recovery. At 12 weeks post nerve injuries, gastrocnemius muscles of both two sides of hind limbs are collected and weighted. LNC and autograft groups display small differences in weight between the damage and normal sides, whereas gastrocnemius muscles exhibit significant atrophy in the sham-operated blank groups, the group involved of photothermal effects alone, and the groups absence of biochemical or bioelectrical cues (Figure 6e, f). Histological analysis via Masson’s staining further demonstrates that gastrocnemius muscles treated by LNC exhibit large-diameter fibers and small percentage of collagen deposition areas, corresponding to improved muscle functions and reduced inflammation, respectively, which are comparable to the autografts and obviously superior over the other experimental groups (Figure 6g, h). Furthermore, we perform compound muscle action potential (CMAP) tests for the hind limbs with injured nerves 12 weeks after surgery and compare them with normal conditions (Figure 6i). The results demonstrate that the LNC-treated nerves can evoke the CMAP amplitude of 1.73 ± 0.03 mV, with no significant differences compared with the contralateral non-treated sites (1.79 ± 0.07 mV, P = 0.137, Figure 6j). However, the CMAP amplitude in the control group displays obvious decreases, implying limited recovery of motor function (Figure 6j). Given the sciatic nerve function index (SFI) can quantitively reflect the recovery of motor functions for sciatic nerve injuries, we also perform gait analysis and record the SFI within the neural regeneration processes (Figure 6k and Figure S19). After postoperative treatment for 4 and 12 weeks, the LNC groups achieve the best efficacy in improving motor functions, as evidenced by the highest SFI scores among all the groups (-78.8 ± 4.2 and -35.7 ± 1.9, respectively), comparable to the autograft-treated conditions (-84.0 ± 1.8, P = 0.163 and -43.2 ± 1.3, P = 0.058, Figure 6l). Overall, compared with existing nerve conduits ranging from conventional topographical constructs to advanced smart platforms^7, 8, 19, 35, 39, 40, 59^, the LNC recapitulates an unexplored biomimetic architecture with coupled physiological signals, achieving exceptional therapeutic efficacy—preventing nerve compression and neuroma formation while promoting robust revascularization and muscle reinnervation. Notably, these outcomes surpass those achieved by the “gold-standard” autograft, offering a transformative avenue to address longstanding challenges in peripheral nerve repair (Figure 6m and Table S1).

## Discussion

In summary, we reported a bioinspired LNC that couples nerve regeneration-like biomechanical, bioelectrical, and biochemical signals, that thereby enabling unprecedented restoration of in-nerve blood supply and reinnervation for motor function recovery for long-distance sciatic nerve amputation. Distinct from autografts and conventional nerve conduits, the LNC features a tri-layered, nerve-like architecture, comprising a hydrogel bilayer, nano-grooved ferroelectric polymer stripes, and living SCs. Leveraging the nerve-like architecture and coupled physiological signals, the LNC achieves exceptional adaptability at the injured nerve interface, enabling nerve tissue-matched electrical conductivity (7.5 ± 0.9 mS·cm^-1^), robust conformal and sutureless tissue integration, and enhanced secretion of pro-regenerative exosomes (3-fold increase; upregulated 42 types of miRNAs related to neural regeneration) and growth factors (2.2- and 1.4-fold enhancements in neurotrophic and angiogenic factors, respectively). When implanted into sciatic nerve of long-distance amputation in rats, the LNC effectively coordinates early immune regulation, in-nerve angiogenesis, accelerated axonal sprouting, and robust remyelination, enabling unprecedented recovery of in-nerve blood supply and motor function (SFI: -35.7 ± 1.9), without suture-induced trauma, tissue compression, neuroma formation, or detectable foreign body responses. We envision that this novel bioinspired and biohybrid design will not only offer a promising translational strategy to overcome longstanding challenges of functional restoration for long-distance nerve injuries but also pave a new way for regenerative medicine and biohybrid brain-machine interfaces^60–63^.

## MATERIALS AND METHODS

### Approval for experiments with animals

All the experiments involving animals were performed under the approval by the Animal Use and Care Committee of the Shenzhen Institute of Advanced Technology, Chinese Academy of Sciences with the project license (SIAT-IACUC-210301-TGS-ZQL-A1667-01) following ethical review.

### Cell culture

Rat Schwann cells (SCs) were obtained from Procell system, which were expanded using Dulbecco’s modified eagle medium (DMEM, Gibco) with supplemented 10 v/v% fetal bovine serum (FBS, ScienCell) and 1 v/v% penicillin/streptomycin solution (P/S, ScienCell), and maintained in an incubator (5 v/v% CO_2_, 37 °C, humid atmosphere, CLM-170B-B-NF, ESCO). After above 70% confluence, SCs were trypsinized and seeded onto sterilized samples at specific cell densities for following investigations.

### Fabrication of bioinspired living nerve conduit (LNC)

LNC was fabricated via stepwise procedures. First, we prepared the ferroelectric polymer striple by compositing poly(vinylidene fluoride-*co*-trifluoro ethylene) (P(VDF-TrFE)) and polydopamine (PDA) via casting, electrospinning, and cutting. Specifically, PDA is synthesized by the polymerization of dopamine hydrochloride (Sigma-Aldrich), which is washed with ethanol, dried, and then dispersed within a solution formulated by dissolving P(VDF-TrFE) (VDF:TrFE molar ratio = 70:30, Mw: 520,000-860,000, Piezotech) in dimethyl sulfoxide (DMSO, Aladdin) with a P(VDF-TrFE) concentration of 10 w/v%. The PDA-to-P(VDF-TrFE) mass ratios were varied from 0.1, 0.5, 1.0, 2.0, to 5.0 wt%. After vigorous stirring, the mixture suspensions were poured into different types of silicon molds, including 2-inch silicon molds with a nanogroove array surfaces (200 nm in depth, 200 nm in width, and 200 nm in edge-to-edge spacing), 6-inch silicon molds with a microgroove array surfaces (3 μm in depth, 10 μm in width, and 10 μm in edge-to-edge spacing), or 6-inch silicon molds with flat surfaces. Followed by drying at 80 °C under ambient pressure overnight and annealing at 130 °C under vacuum for 6 h, PDA/P(VDF-TrFE) films were peeled off from the silicon molds, and subsequently treated with corona polarization at 22 kV for 25 min. For comparative studies, the films made of PDA and non-ferroelectric poly(vinyl chloride) (PVC, Mw: ∼233,000, Sigma-Aldrich) were also fabricated using the above-describe methods. Afterwards, the poled PDA/P(VDF-TrFE) films were deposited with a layer of electrospun PDA/P(VDF-TrFE) fibers, which were formed by electrospinning of a precursor formulated by dissolving 20 w/v% P(VDF-TrFE) in *N*,*N*-dimethylformamide (DMF, Yonghua Fine Chemicals Co., Ltd.) and acetone with a volume ratio of 6:4. The electrospinning was performed at 15 kV for 5 h, with a working distance of 10 cm and a feeding speed of 1 mL·h ^-1^. Subsequently, these composite films were cut into stripe arrays (with a width of 0.5 mm, an edge-to-edge distance of 0.5 mm, and a length of 15 mm) by using a pre-machined mold, resulting in the ferroelectric polymer stripes. Second, the hydrogel bilayer, i.e., calcium-ion-crosslinked, *N*-hydroxysuccinimide-functionalized alginate and bio-ionic liquid-conjugated poly(acrylic acid) (Alg-NHS/PAAc-IL) hydrogel and chitosan hydrogel, were sequentially integrated with the ferroelectric polymer stripes via thermo-polymerization and ionic crosslinking. Typically, Alg-NHS/PAAc-IL was prepared by dissolving 1.5000 g acrylic acid (Sigma-Aldrich) with an equimolar ratio of choline bicarbonate (Sigma-Aldrich) in 10 mL of water, followed by the addition of 5.0000 g sodium alginate (Sigma-Aldrich), 0.1450 g *N*-hydroxysuccinimide (Sigma-Aldrich), 0.2425 g 1-ethyl-3-(3-dimethylaminopropyl) carbodiimide (Sigma-Aldrich), 0.0600 g ammonium persulfate (Sigma-Aldrich), and 0.1200 g poly(ethylene glycol) diacrylate (Mn: 700, Sigma-Aldrich), which were subsequently cross-linked within a rectangular confined cell made of two glass plates and a 100-μm poly(tetrafluoroethylene) spacer, and then onto the ferroelectric polymer stripes for another 4 h. After drying the Alg-NHS/PAAc-IL at room temperature in air, a chitosan (Sigma-Aldrich) solution (2 w/v% in 2 v/v% acetic acid-water solvent) was poured onto the films and then dried in air at 37 °C and 60% humidity for over 8 h for evaporating the solvents, immersed in a 0.4 M CaCl_2_ solution and phosphorate buffer saline (PBS, Sigma-Aldrich) for ionic crosslinking until swelling equilibrium, obtaining the hydrogel-PDA/P(VDF-TrFE) composite films. Third and finally, sterilized composite films were seeded with SCs with specific cell densities, therefore resulting in LNC.

### Characterizations

The macroscopic shapes of samples, as well as their self-rolling processes, were recorded by a digital camera (EOS 6D Mark II, Canon). To characterize the microscopic structures and morphologies, samples with a thin layer of Au sputtering were observed under a field-emission scanning electron microscope (SEM, Sigma 300, Zeiss), both at the top and cross-sectional views.

To characterize the compositions of LNC, we performed energy dispersive spectroscopy to map the elemental distribution of N (for identifying PDA), F (for identifying P(VDF-TrFE)), and Ca (for identifying alginate-based hydrogel). We also performed Fourier transform infrared spectroscopy (FTIR, Smartlab 9, Rigaku) to examine the fraction of ferroelectric β phase P(VDF-TrFE). According to Beer-Lambert equation, the fraction of ferroelectric β phase P(VDF-TrFE) (F(β)) was calculated according to the following equation (1):

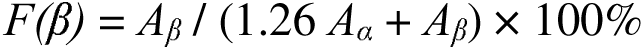

where *A*_α_ and *A*_β_ represent the characteristic absorption peaks to the α-phase (764 cm^-1^) and β-phase (843 cm^-1^) of the P(VDF-TrFE), respectively.

To characterize the mechanical properties of LNC, we performed tensile tests for the samples on a universal testing machine (Hengyi Precision Instrument Co., Ltd.). The strain-stress curves were recorded and their mechanical properties such as ultimate tensile strength, fracture strain, and Young’s modulus, were then calculated accordingly. In addition, we also quantified the shear strength for the bio-adhesive interface between the LNC and *ex vivo* rat sciatic nerves by means of lap shear test using the universal testing machine. Two fresh nerve segments were enwrapped by the self-rolled LNC, which were then clamped separately by the machine to record the force and displacement under tensile. The shear strength for the bio-adhesive interface was calculated according to the following equation (2):

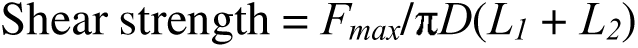

where *F_max_*represents the maximum force, *D* represents the outer diameter of the nerve segments, *L_1_* and *L_2_* represent the overlapping lengths between the hydrogel and nerve segments.

To characterize the electrical properties of LNC, we measured the ionic conductivity for the hydrogel bilayer of LNC by means of AC perturbation. In brief, the hydrogels were clamped between two ITO conductive glasses. The impedance of the sample was then recorded by using a LCR digital bridge (Tonghui Electronic) by applying 1 V AC at 10 kHz. The conductivity (σ) was calculated according to the following equation (3):

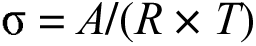

where *A* represents the area of the glasses covered by the hydrogel, *R* represents the impedance, and *T* represents the thickness of the hydrogel.

We also measured the open-circuit voltages generated by the ferroelectric polymer layers of LNC under near-infrared (NIR) irradiation for evaluating their photo-pyroelectric properties. In brief, we monitored surface temperature of the samples under 808 nm NIR irradiation (spot area of 1 cm^2^, pulse width: 500 ms, frequency: 1 Hz) generated by a NIR laser with a built-in digital shutter (Beijing Hong Optoelectric Co. Ltd.) using a high-resolution infrared thermometer (R550Pro, 21 μm, NEC). Afterwards, we then record the open-circuit voltages generated by the ferroelectric polymer layers using a digital source meter (2470, Keithley). In the recording, 200-nm ITO was coated onto both sides of the samples (cut into square shapes with a width to length of 1.20 cm × 1.20 cm) as transparent electrodes. The photo-pyroelectric properties of the samples with different PDA contents (0.1, 0.5, 1.0, 2.0, and 5.0 wt%), under various power densities (5, 10, 20, 30, and 40 mW·mm^-2^), and treated with 37 °C PBS immersion for different time periods (4, 8, and 12 weeks) were evaluated using this methods. Moreover, we measured the current generated by the ferroelectric layer of the LNC using a patch-clamp setup. Briefly, we immersed the samples in electrolyte-contained phosphorate buffer saline (PBS, Gibco, pH: 7.4) to simulate physiological condition. An Axopatch 200B amplifier (Molecular Devices) with a Ag/AgCl electrode was used in the measurements at the voltage-clamp mode, where the glass pipettes filled with PBS was placed near the smaple (∼5 μm). Data was acquired at room temperature under the NIR irradiation (power density: 5 mW·mm^-2^) towards the samples and analyzed using the pClamp software (Molecular Devices). The photo-pyroelectric properties of the ferroelectric polymer layers of LNC after 12-week implantation in a rodent model were further assessed through scanning Kelvin probe microscopy (SKPM, Asylum Research Cypher S). Surface potential was measured for the samples coated with a grounded silver paste under 808 nm near-infrared laser irradiation at 2.5 min intervals at room temperature.

### Cell viability and proliferation assessment

The viability of SCs within LNC was evaluated through staining them using a LIVE/DEAD Viability/Cytotoxicity Kit (Invitrogen) and visualization under a laser scanning confocal microscope (LSCM, Zeiss, LSM 900). SCs were seeded on sterilized samples in a 24-well plate at the density of 1,000 cells·cm^-2^. After 24-h incubation, SC-laden samples were treated daily by NIR irradiation (1 Hz, 500 ms, spot area of 1 cm², 2-min irradiation with 2-min interval each turn, 5 turns per day) with various power densities (10, 20, 30, and 40 mW·mm^-2^) for different time periods (1, 4, and 6 days). Afterwards, cells were stained with the LIVE/DEAD Viability/Cytotoxicity Kit, where live and dead cells were labelled by Calcein AM (in green) and ethidium homodimer-1 (in red), respectively. Percentage and density of live cells were analysed from the LSCM images with the assistance of the ImageJ software.

### Cell migration assessment

To assess cell migration on the ferroelectric polymer samples with different types of surface morphologies (including nano-groove array surfaces, micro-groove array surfaces, and flat surfaces), we cultured SCs on the sterilized samples at the density of 40,000 cells·cm^-2^ with the assistance of a two-channel mold (length: 5 mm, channel width: 2 mm, channel distance: 500 μm), where the orientation of the channels were parallel to the orientation of nano-/micro-groove arrays. After 24-h incubation for SC attachment, the molds were removed. Cell patterns at the initial status immediately after the removal of the mold were recorded by using an inverse microscope (Olympus X71). 12 h later, the patterns of SCs cultured on the different types of surfaces were examined by the inverted microscope again. Cell migration velocity was analyzed according to the bright-field image (N = 5) captured by the microscope with the assistance of the ImageJ software.

### Intracellular Ca^2+^ imaging

To examine whether the bioelectrical signals generated by the ferroelectric polymer layers of LNC can elicit Ca^2+^ signalling of the incorporated SCs, we cultured SCs onto the sterilized PDA/P(VDF-TrFE) or PDA/PVC samples in a 96-well plate at the density of 15,000 cells·cm^-2^. After 48-h incubation, SC-laden samples were washed with warm PBS one time, incubated with pre-diluted Fluo-4 AM (Invitrogen) at a dilution of 1/100 in DMEM according to the manufacture’s protocol in dark at 37 °C for 30 min, and then incubated with warm artificial cerebrospinal fluid (aCSF; containing 126 mM NaCl, 2.5 mM KCl, 2 mM CaCl, 2 mM MgCl, 1.25 mM NaH PO, 26 mM NaHCO, and 10 mM glucose, pH 7.4) in dark at 37 °C for another 30 min. Subsequently, time-lapse fluorescence images were recorded using the LSCM for 30 s before, 60 s during, and 60 s after NIR irradiation (20 mW·mm^-2^, 1 Hz, 500 ms, spot area: 1 cm^2^). The fluorescence intensity of each cell was measured using ImageJ. The normalized fluorescence changes (ΔF/F_0_) were calculated according to the following equation (4):

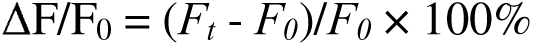

where *F_0_*represents the initial fluorescence intensity during recording, *F_t_*represents the real-time fluorescence intensity during recording, Δ*F* represents the change in the fluorescence intensity.

### Exosome characterizations and quantification

To quantify the exosome secreted by LNC, we first cultured SCs on the ferroelectric polymer samples with different types of surface morphologies, including nano-groove array surfaces, micro-groove array surfaces, and flat surfaces. After 24 hours for SC attachment, we changed the medium to the exosome-free medium (DMEM with supplemented 10 v/v% Exosome-depleted FBS (Dakewe Biotech Co., Ltd.) and 1 v/v% P/S). The medium for the samples after specific time periods and/or specific NIR irradiation treatments (20 mW·mm^-2^, 1 Hz, 500 ms, spot area of 1 cm², 2-min irradiation with 2-min interval each turn, 5 turns per day) were then collected and centrifuged at 2,000 g, 4 °C for 30 min to remove debris. The supernatants were then mixed with a Total Exosome Isolation Kit (from cell culture media, Invitrogen) according to manufacturer’s protocol, incubated at 4 °C overnight, centrifuged at 10,000 g, 4 °C for 1 h, which were subsequently resuspended with PBS and stored at -20 °C for further characterizations.

The exosomes derived from different types of samples were characterized by transmission electron microscopy (TEM), western blot (WB), and nanoparticle tracking analysis (NTA) for identification and quantification. To be specific, the morphologies of exosomes were characterized by using a transmission electron microscope (HT7800, HITACHI). The samples for TEM characterization were negatively stained with 12-wolframophosphoric acid (RHAWN Co., Ltd.) and air dried at room temperature. The characteristic proteins to exosomes, including TSG101, CD63, and Hsp-70, were characterized via WB, by using GAPDH as internal reference. In the WB assay, exosome-suspended samples were treated with cold RIPA lysis buffer (containing 1× protease inhibitor cocktail and 1 mM phenylmethanesulfonyl fluoride), oscillated on ice for 30 min, and then centrifuged at 14,000 rpm, 4 °C for 15 min. The supernatant was then collected and boiled at 100 °C for 5 min to extract protein. Protein concentration was examined using a BCA assay kit (Thermo Scientific), and samples were adjusted to uniform concentration. Proteins were separated by SDS-PAGS under constant voltage and transferred to a methanol-activated PVDF membrane. This membrane was further blocked with 5% defatted milk in Tris-borate-sodium Tween-20 buffer (TBST) at room temperature for 2 h, incubated with primary antibodies [rabbit polyclonal anti-GAPDH antibody (Abcam, ab9485), rabbit monoclonal anti-TSG101 antibody (Abcam, ab133586), rabbit polyclonal anti-CD63 antibody (Abcam, ab217345), and Mouse monoclonal anti-Hsp70 antibody (Abcam, ab5439)] at 4 °C overnight, and then incubated with an HRP-conjugated secondary antibody at room temperature for 2 h. Patterns on the membrane were captured using a gel imaging system. To quantify the exosomes, we collected the exosome from different groups, resuspended them in 1 mL PBS, and then analyzed the size distribution and quantities of exosomes using a NTA instrument (NS300, NanoSight) Absolute number of exosomes within the diameter range of 0-200 nm and the amounts of SCs were quantified, which were used to calculate the number of exosome secreted per cell in each group.

### Bulk small RNA-seq

To sequence the bulk small RNA within exosomes secreted by SCs within LNC, we cultured the SCs and collected exosomes according to the above-described procedures. We then lysed the exosomes with Trizol (Invitrogen). Afterwards, the exosomes samples were mixed with cold trichloromethane (TCM, Yonghua Fine Chemicals Co., Ltd.) at 4 °C for 2 h, centrifugedat 14,000 rpm, 4 °C for 15 min. The supernatant was then collected, mixed with cold isoproryl alcohol (IPA, Dongguan Dongjiang Chemical Reagent Co., Ltd.) at 4 °C for 20 min, and centrifuged at 14,000 rpm, 4 °C for 15 min. The precipitation was collected, resuspended with 75% ethanol configured with diethyl pyrocarbonate (DEPC)-treated water, centrifuged at 14,000 rpm, 4 °C for 15 min, and dissolved in DEPC-treated water to obtain small RNA samples. After establishing library using QIAseq miRNA Library Kit (QIAGEN), those small RNA samples were sequenced and analyzed using NextSeq CN500 (Berry Genomics Co., Ltd.) and Fast-QC software, respectively. Subsequently, the Gene ontology (GO) enrichment analysis and Kyoto encyclopedia of genes and genomes (KEGG) enrichment analysis were performed via miEAA, a web-based tool for functional analysis of sets of miRNAs, refer to miEAA 2023: updates, new functional microRNA sets and improved enrichment visualizations.

### Enzyme-linked immunosorbent assay (ELISA)

To examine the growth factor secretion, we performed ELISA for the supernatant collected from different experimental groups. Specifically, we cultured SCs on sterilized samples in a 24-well plate at the density of 2,500 cells·cm^-2^. After 24-h incubation, the SC-laden samples were treated with the chosen NIR irradiation mentioned above on the following 1^st^ and 4^th^ days. Concurrently, the supernatant of each sample was collected daily, centrifuged at 2,000 rpm, 4 °C for 30 min, collected the supernatants, and then stored at -80 °C prior to the ELISA measurements. After collection of all the supernatants, the concentrations of nerve growth factor (NGF) and vascular endothelial growth factor (VEGF) in these supernatants were measured using rat NGF beta/NGF ELISA kit (Boster) and rat VEGF standard ABTS ELISA development kit (PEPRO TECH, 900-K436) following the manufacture’s protocols, which were quantified according to the absorption at 405/650 nm using a microplate reader (Molecular Devices, Cmax Plus) and their respective calibration curves obtained from standard solutions with pre-determined gradient concentrations.

### Transcriptome analyses

Bulk RNA sequencing was performed for the SCs within LNC to investigate the underlying mechanisms how biomechanical and bioelectrical signals modulate SC behaviours. SCs for bulk RNA sequencing were seeded on sterilized samples in a 24-well plate at the density of 3,250 cells·cm^-2^. After 24-h incubation, SC-seeded samples were treated daily using the chosen NIR irradiation mentioned above for the next 3 days. After 3-day treatment, these samples were lysed with Trizol for extracting mRNA through the above-mentioned procedures. Subsequently, the concentrations and purities of mRNA samples were examined, followed by reverse transcription of mRNA into cDNA. After establishing library, cDNA was sequenced using Illumina PE150 and analyzed using the StringTie software. GO enrichment analysis and KEGG enrichment analysis were subsequently performed via clusterProfiler software.

### Transwell experiments

To evaluate the pro-regenerative potential of LNC in vitro, we co-cultured LNC, as well as control samples including blank control (SC-free samples), with different types of cells in a transwell setup (Nest), which contains separate chambers for LNCs and the co-cultured cells, divided by a network that allow for inter-diffusion of biochemical signals yet impede intercellular contact. For the transwell experiments, we first seeded SCs onto sterilized samples at a cell density of 2,000 cells·cm^-2^ in 96-well plate. 24 hours later for SC attachment, we then transferred the SC-laden samples into the upper chambers, where the bottom chambers were pre-seeded with rat-derived macrophage (Procell), phaeochromocytoma cells (PC12, Procell, as neuronal model) or human umbilical vein endothelial cell line (Ea.hy926, Procell, as endothelial cell model). After 6-h coculturing in the transwell setup, NIR irradiation (15 mW·mm^-2^, 1 Hz, 500 ms, spot area of 1 cm^2^, 2-min irradiation with 2-min interval each turn, 5 turns per day) was exploit during the next 3 days. To assess the pro-angiogenic potentials, the Ea.hy926 cells were seeded onto the bottom chamber with pre-coated a thin layer of Matrigel (Corning) at a cell seeding density of 94,000 cells·cm^-2^. The formation of vascular network for each sample was examined under a bright-field microscope (Ti-U, Nikon). The numbers of rings and nodes for each sample were accounted and statistically analyzed according to the captured bright-field images with the assistance of the ImageJ software. Additionally, SCs in the transwell experiments were stained with PKH26 (Sigma) according to the manufacture’s protocol. After 3-day coculture, PC12 cells were washed with PBS three times and then visualized under the LSCM for tracking the uptake of SC-secreted exosomes by neighbouring cells.

### Reverse transcription quantitative polymerase chain reaction (RT-qPCR)

To quantify the expression of Tuj1, we also performed RT-qPCR for the cocultured PC12 cells. Specifically, PC12 cells after coculturing were washed with PBS three times, lysed with Trizol, and then extracted with trichloromethane-phenol-ethanol. After quality control. RT-qPCR was performed using Hifair® 1st strand cDNA synthesis SuperMix for qPCR (gDNA digester plus) (Yeasen Biotechnology (Shanghai) Co., Ltd., 11141ES60) on PCR amplifier (Bio-Rad), followed by Hieff UNICON® Universal Blue qPCR SYBR Green Master Mix (Yeasen Biotechnology (Shanghai) Co., Ltd., 11184ES08) on LightCycler 96 (Roche). All experiments were replicated three times. The primer sequences designed in the RT-qPCR analysis are listed in Table S1.

### Animal model, surgery, and treatment

Male Sprague-Dawley rats (8∼12 weeks old) were used to establish the rodent model with long-distance sciatic nerve injuries. To establish the model, we anesthetized the rats via intraperitoneal injection of pentobarbital sodium (1 w/v%) at a dosage of 80 mg/100 g. An incision was then created at the right hindlimb, exposing sciatic nerves. a10-mm defect between the proximal and the distal nerve bundle was then created by cutting off the middle segment of the exposed sciatic nerve. The rat model with such long-distance sciatic nerve injures was then treated by different methods/samples, which could be divided into five groups: i) Blank: subjects with the incision being sutured, without any implants on the defect sites, followed by NIR irradiation treatment since post-surgery day 3; ii) Autograft: subjects with the resected nerves reversely sutured onto the defect sites, followed by NIR irradiation treatment since post-surgery day 3; iii) Control: subjects with the defect sites implanted with PVC-based samples (i.e., tubes consisted of hydrogel bilayer and PDA/PVC stripes), followed by NIR irradiation treatment since post-surgery day 3; iv) SC-free: subjects with the defect sites implanted with SC-free samples (i.e., tubes consisted of hydrogel bilayer and PDA/P(VDF-TrFE)), followed by NIR irradiation treatment since post-surgery day 3; v) LNC (NIR-): subjects with the defect sites implanted with LNC, without NIR irradiation after the surgery; vi) LNC (NIR+): subjects with the defect sites implanted with LNC, followed by NIR irradiation treatment since post-surgery day 3. After the surgery, all the rats were injected penicillin per day for the next 3 days to prevent infection. NIR irradiation was performed using the parameters of 15 mW·mm^-2^, 1 Hz, 500 ms, spot area of 1 cm^2^, 2-min irradiation with 2-min interval each turn, and 5 turns per day. To measure localized temperature changes of tissues under the NIR irradiation, we co-implanted a thermocouple thermometer and monitored the temperature changes using a multichannel signal acquisition system (TB-9212, National Instruments).

### *In vivo* SC tracking

To track the long-term bioavailability and viability of SCs within LNC after in vivo implantation, we transfected green fluorescent protein (GFP) AAV packaging plasmids (Shanghai HanBio Co., Ltd.) into the SCs and prepared the LNC for implantation using these GFP-expressed SCs. After 4- and 12-week implantation, the implanted LNC were collected and observed under the LSCM.

### Micro-CT imaging

We performed micro-CT imaging for the hindlimbs of subjects with the implants, with the sciatic nerve tissues stained with iohexol. The samples were scanned using a high-resolution micro CT system (SkyScan 1272, Shuyun Instrument) and reconstructed using the NRecon software. 3D visualization with pseudo-color was carried out using the CTvox software.

### Photoacoustic imaging

We also performed photoacoustic imaging for normal rats (without injuries), as well as subjects in the LNC (NIR-) and LNC (NIR+) groups. The animals for photoacoustic imaging were anesthetized with isoflurane. Their skins and muscular tissues were then incised to expose the sciatic nerve with or without implants. The exposed nerves were then scanned and visualized under a lab-made photoacoustic imaging system, where line scanning for the vessel in the region of interest was conducted for quantifying the flow rate of blood.

### Gastrocnemius muscle assessment

To evaluate post-operative muscle atrophy, wet weight the isolated gastrocnemius muscles for both hindlimbs of subjects in the five groups. The ratio of the injured side with respective to the normal side for each subject was calculated. In addition, we also performed Masson’s staining for the isolated gastrocnemius muscles, which were crossly sectioned into slices by a cryostat microtome (CM1860, Leica) and stained by using a Masson’s staining kit (Servicebio). The stained slices were visualized under a stereomicroscope (SZ61, Olympus). the percentages of collagen area and the diameter of muscle fibres were calculated according to the bright-field images with the assistance of the ImageJ software.

### Electrophysiological assessment

To evaluate the recovery of motor functions for the subjects under different types of treatment, we record the compound muscle action potentials (CAMP) for the rats at post-operative 12 weeks using a portable biofunctional experimental system (BL-420S, Taimeng, China). The CAMP under the periodic alternating electrical stimulation (25 mV, 25 ms, 0.1 Hz) were recorded, where their peak amplitudes were then calculated.

### Gait analyses

The recovery of motor functions for the subjects under different types of treatment for 4 and 12 weeks was also evaluated through gait analyses using a CatWalk-automated gait analysis system (XR-FP101, Shanghai Xinruan Information Technology Co. Ltd., China) with the assistance of a built-in software. The subjects were pre-trained twice. Then they were allowed to pass through a glass walkway of the system, where their footprints were recorded by a high-speed camera mounted under the glass. Sciatic function index (SFI) was calculated according to the footprints using the following equation (5):

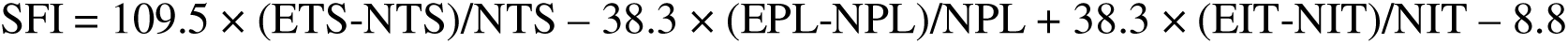

where E and N represent the experimental and normal sides, respectively, TS represents the toe spread (distance between the first and fifth toes), PL represents paw length (distance between the third toe and the heel), IT represents intermediary toe spread (distance between the second and fourth toes). The SFI ranges from -100 to 0, with -100 indicating complete disfunction and 0 indicating normal function.

### Histological assessment

At post-operative 12 weeks, we collected sciatic nerve tissues at the implanted sites and then treated them with hematoxylin and eosin (H&E) staining for histological assessment. The collected tissues were fixed in cool 4 v/v% PFA, dehydrated with an ascending ethanol gradient, embedded in paraffin, and then longitudinal sectioned into slices using the cryostat microtome. Subsequently, the slices were stained with H&E HD constant dye kit (Servicebio) according to manufacturer’s protocol, rehydrated with a descending ethanol gradient, and then mounted with neutral balsam (Sinopharm Chemical Reagent Co., Ltd.). Finally, the images of the slices were observed and captured under the stereomicroscope.

### Immunofluorescence assessment

We first performed immunochemical staining for the SCs cultured on different types of samples for examining the influences of surface morphologies on cellular shapes. To be specific, SCs were seeded on sterilized samples with flat, micro-groove array, or nanogroove array morphologies at a density of 4,000 cells·cm^-2^. After incubation for 96 h, these samples were washed with PBS three times, fixed with 4 v/v% paraformaldehyde (PFA, Sigma-Aldrich)/PBS solution for 10 min, permeated and blocked with PBS with supplemented 0.1 v/v% Tween-20 (Sigma) and 1 w/v% bovine serum albumin (BSA, Sigma) for 2 h, incubated with S100 polyclonal antibody (Invitrogen) at a dilution of 1/1000 in PBS with supplemented 0.1 v/v% Tween-20, 1 w/v% BSA, 10 v/v% goat serum (Sigma-Aldrich), and 0.3 M glycine (Sigma-Aldrich) at 4 °C overnight, and then stained with Goat anti-rabbit IgG H&L (Alexa Fluor® 568) (Abcam, ab175471) and 4’6-diamidino-2-phenylindole (DAPI, Abcam, Ab104139) at a dilution of 1/1000 in PBS with supplemented 1 w/v% BSA for 1 h for labeling S100 protein and nucleus, respectively. Subsequently, fluorescence images for these stained samples were captured by using the LSCM. Cellular orientations on the samples with different types of morphologies were analyzed through measuring the angles of long axis of cell shapes with respect to the y-axis (defined by the orientation of micro-/nano-groove arrays) according to the LSCM images with the assistance of the ImageJ software.

We also performed immunochemical staining for the cocultured rat-derived macrophages and PC12 cells in the transwell experiments for examining the phenotypes and the expression of beta III Tubulin (Tuj1). Briefly, co-cultured macrophages and PC12 cells were fixed, permeated, and blocked following the above-described procedures. Afterwards, macrophages were incubated with anti-CD206 rabbit pAb (Servicebio) and goat anti-mouse IgG H&L (Alexa Fluor® 555, Abcam, ab150114), followed by incubation with recombinant anti-iNOS antibody (rabbit mAb, Servicebio) and goat anti-mouse IgG H&L (Alexa Fluor® 488, Abcam, ab150113). PC12 cells were then incubated with recombinant rabbit monoclonal to Tuj1 (Abcam, ab52623) and goat antirabbit IgG H&L (Alexa Fluor® 488) (Abcam, ab150077). These stained cell samplers were visualized under the LSCM. The relative fluorescence intensity of iNOS and CD206 for the stained macrophages, as well as the length-to-width ratios and relative fluorescence intensity for the PC12 cells with stained Tuj1 were analysed according to the LSCM images with the ImageJ software.

In the *in vivo* investigations, we collected sciatic nerve tissues at the implanted sites at post-operative 12 weeks and also performed immunofluorescence assessment on them. For immunostaining, tissues at nearly half-of-middle were embedded in optimal cutting temperature compound, which were then crossed sectioned into slices using the cryostat microtome. Subsequently, the slices were first blocked with 3 v/v% methanol (Sinopharm Chemical Reagent Co., Ltd.) in hydrogen peroxide 30% water solution (Sinopharm Chemical Reagent Co., Ltd.) at room temperature for 25 min, second blocked with 3 w/v% BSA in PBS at room temperature for 30 min. Thereafter, the slices were successively incubated primary antibodies at 4 °C overnight and corresponding second antibodies at room temperature for 50 min. To evaluate the ratios between M1 and M2 type macrophages, the treated slices were stained with successively incubated anti-CD206 rabbit pAb (Servicebio), recombinant anti-iNOS antibody (rabbit mAb, Servicebio), and then goat anti-mouse IgG H&L (Alexa Fluor® 555, Abcam, ab150114, for CD206) and goat anti-mouse IgG H&L (Alexa Fluor® 488) (Abcam, ab150113, for iNOS), respectively. To evaluate the densities of nerve fibers, myelin sheathes, and vessels, the slices were stained with anti-neurofilament heavy polypeptide mouse mAb (NF200, Servicebio), recombinant anti-S100 beta antibody (mouse mAb, Servicebio), anti-CD31 rabbit pAb (Servicebio), and then goat anti-mouse IgG H&L (Alexa Fluor® 488) (Abcam, ab150113, for NF200), goat anti-mouse IgG H&L (Alexa Fluor® 555) (Abcam, ab150114, for S100), and goat anti-mouse IgG H&L (Alexa Fluor® 647) (Abcam, ab150115, for CD31), respectively. Nuclei were stained with DAPI. The stained slices were observed under the LSCM. The area fraction for each biomarker was analyzed according to the fluorescence images with assistance of the ImageJ software.

### Transmission electron microscopy

The regenerated tissues collected at post-operative 12 weeks were further observed under a a transmission electron microscope (TEM, HT7800, Hitachi) for evaluating the thickness of myelin sheath and the diameter of nerve fibers. Tissues for TEM observation were fixed in electron microscope fixative (Servicebio) in dark at 4 °C, post-fixed with a OsO_4_ solution (0.1 M in PBS, Ted Pella Inc.) for 2 h at room temperature, dehydrated with an ascending ethanol gradient, embedded in EMBed 812 at 37 °C overnight, polymerized at 60 °C for 48 h, and then crossed sectioned into slices. After these treatments, the slices were successively stained with 2 w/v% uranium acetate (in saturated ethanol) solution at dark for 8 min and then 2.6 w/v% lead citrate (Sigma) for 8 min, followed by the TEM observation. The thickness of myelin sheath and the diameter of nerve fibers were measured according to the TEM images.

### Numerical modeling

Using open-source codes mcxyz.c (https://omlc.org/software/mc/mcxyz/index.html), we performed numerical modeling according to the established Monte Carlo modeling of light transport in tissues to simulate the NIR penetration through epidermis (thickness: 2 mm) and dermis (thickness: 8 mm), equivalent to the conditions the transdermal irradiation towards peripheral nerves. The parameters of NIR light irradiated to the tissues were set with wavelength of 808 nm, spot size of 10 mm, and power density of 2 W cm^-2^, which are used in our *in vivo* investigations. The output files were convolved and illustrated through the software MATLAB.

### Statistical analysis

Data are presented as means ± standard error of mean (S. E. M.). The numbers (n) of biological replicates or mice are indicated in corresponding figure legends. We performed statistical analysis using one-way analysis of variance (ANOVA) followed by the two-tailed Student’s *t*-test, as indicated in corresponding figure legends.

## Acknowledgments

The authors acknowledge the financial support provided by the National Natural Science Foundation of China (52261160380, 52022102, 52173148), the Shenzhen Medical Research Fund (B2302045), Research Grants Council of Hong Kong (N_HKU749/22), the Youth Innovation Promotion Association of CAS (Y2023100, 2022368), the Fundamental Research Program of Shenzhen (RCJC20221008092729033, JCYJ20220818101800001).

## Author contributions

X. D conceived the idea and supervised the research. Y. C. and Q. Z. conducted the experiments with the help of photoacoustic imaging by X. L. and C. L.. Q. Z. and X. D analyzed the results and wrote the manuscript with the assistance of G. L., M. W. and Y. Y.. All authors contributed to the discussion and interpretation of the results. Y. C, and Q. Z. contributed equally to this work

## Data, Code, and Materials Availability

All data and code needed to evaluate and reproduce the results in the paper are present in the paper and/or the Supplementary Materials. This study did not generate new materials. All chemicals and reagents used are commercially available, and their sources are specified in the Materials and Methods section.

**Supplementary Materials** Supplementary Figures S1 to S19 Supplementary Tables S1 to S2 Supplementary Videos 1 to 4

